# Phenotypic plasticity, population structure and adaptation in a young weed species with a worldwide distribution

**DOI:** 10.1101/2020.11.27.401562

**Authors:** A. Cornille, M. Tiret, A. Salcedo, H.R. Huang, M. Orsucci, P. Milesi, D. Kryvokhyzha, K. Holm, X.J. Ge, J.R. Stinchcombe, S. Glémin, S.I. Wright, M Lascoux

## Abstract

The colonization success of a species depends on phenotypic plasticity, adaptive potential and population structure. Assessing their relative contributions during a colonization process is challenging, and a large-scale experiment had yet to be done. In this study, we attempted to tease apart their effects on the fitness of one of the most common plant on Earth, the shepherd’s purse (*Capsella bursa-pastoris*), a self-fertilizing and allopolyploid weed, with a worldwide distribution. The overarching goal is to eventually understand how the shepherd’s purse extensive distribution range was established so rapidly. To do so, we carried out three common gardens, located in Europe, Asia and North America, and measured several life-history traits on field-collected accessions belonging to three distinct genetic clusters (Middle East, Europe, and Asia). Our experiment showed that (i) the success of *C. bursa-pastoris* is mainly due to its high degree of phenotypic plasticity; and (ii), genetic cluster effect reflected a classic pattern observed in core *vs* marginal populations, with the Middle Eastern cluster (putative core population) outperforming the European and Asian clusters. This study therefore revealed, in a model species, different relative contributions of plasticity and adaptation to fitness, depending on the population and the time since colonization occurred.

## Introduction

Some species have tiny natural ranges, others very large ones. In the latter, the current size was sometimes reached surprisingly fast something that might have left a strong footprint on the current pattern of genetic and phenotypic variation. In the present study we will focus on such a case, namely the shepherd’s purse, *Capsella bursa-pastoris* (L.) Medik. (*Brassicaceae*), one of the most common plants on Earth (Coquillat, 1951). Since plants are sessile organisms, the expansion speed mainly depends on their ability to disperse efficiently and establish themselves successfully in new environments. Sometimes the environment will be quite similar to the one in which the parental plants were growing, sometimes it will not. In this latter case, establishment success in the short term may depend on phenotypic plasticity and in the long run on the capacity to adapt to local conditions. Phenotypic plasticity is defined here as the ability of a given genotype to produce different phenotypes in different environments (Bradshaw, 1965; Grenier *et al*. 2016 and references therein), and adaptation as the natural selection process by which a population progressively increases its fitness in a given environment (Linhart & Grant, 1996; Chevin *et al*. 2010 and references therein). These two evolutionary mechanisms are not mutually exclusive, so that each successful expansion will likely correspond to a particular combination of both. In any case, when species succeed to extend their distributions to a worldwide scale, granted that they hence face strongly different environmental conditions, signatures of both adaptation and phenotypic plasticity are likely to be observed. We can conjecture that phenotypic plasticity might be more important in newly established populations and adaptation in older ones. Such predictions are fraught with difficulties, since the age of populations is generally hard to estimate and patterns of phenotypic plasticity may vary depending on the time since colonization. Indeed, theoretical models indicate that while adaptation to a new extreme environment may lead to a transient increase in plasticity, this is followed by a second period of genetic assimilation which, on the contrary is associated to decreased plasticity (Lande, 2015). These complex dynamics could explain why different studies on phenotypic plasticity of colonizing species led to divergent conclusion: some recent studies concluded that phenotypic plasticity could play a role during colonization of new environments (e.g., Daehler, 2003) while others did not (e.g., Davidson *et al*. 2011, Godoy *et al*. 2011). Likewise, while evolutionary pressures underlying adaptation in populations at equilibrium are fairly well documented, the literature is more limited for recent and marginal populations that are usually characterized by non-equilibrium demographics (Excoffier *et al*., 2009; Gilbert *et al*., 2017). As illustrated by Willi *et al*. (2018) determining the respective role of phenotypic plasticity and adaptation during colonization can be challenging.

Two prominent properties of plants that might influence a plant’s adaptive potential and its phenotypic plasticity are ploidy level and mating system. Both have their own costs and benefits, though arising on a different timescale. Allopolyploidy and its evolutionary success can appear paradoxical since its birth will also be accompanied by numerous early challenges (Yant & Bomblies, 2015; Pelé *et al*., 2018). These challenges are first encountered during the initial hybridization event between two divergent genomes, implying, among other things, potential changes of gene expression patterns (Bomblies *et al*., 2016). On the other hand, the increase in chromosome number in polyploids creates genetic redundancy, which potentially allows pattern of expression to diverge, and the evolution of new functions as well as tissue-specific expression of different gene copies (Buggs *et al*., 2010, 2011). This greater genomic flexibility could lie behind the increased trait plasticity of tetraploids in heterogeneous environments, as was recently observed in strawberries (Wei *et al*., 2019). As to mating system, the shift from out-crossing to self-fertilization (a.k.a. selfing) confers “reproductive assurance” when the number of mates is limited. Reproductive assurance is expected to favor colonization of new environments as a few individuals can establish a new population (Baker *et al*. 1965; Pannell *et al*., 2015). The benefit of self-fertilization might, however, be short-lived because the lack of genetic mixing is predicted to limit adaptation and to lead to the genome-wide accumulation of deleterious mutations (Heller & Smith, 1978; Hollister *et al*., 2015, Glémin *et al*., 2019). In agreement with these predictions, selfing species tend to have larger ecological ranges (Grossenbacher *et al*., 2015) but decreasing niche breadth with time (Park *et al*., 2018) compared to their outcrossing congeners, aligned with the “evolutionary dead end” hypothesis (Stebbins 1957; Takeyabashi & Morell, 2001 and references therein). This phenomenon is even more pronounced during a colonization process (Slatkin & Excoffier, 2012; Gonzalez-Martinez *et al*., 2017 and references therein).

*C. bursa-pastoris* is a successful worldwide self-fertilizing colonizer of recent allopolyploid origin that arose some 100,000 years ago from the hybridization between the self-fertilizing, *C. orientalis* (Fauché & Chaub.) Boiss. and the out-crossing, *C. grandiflora* (Klokov) (Douglas *et al*., 2015). As opposed to its two parents, which are restricted to specific areas (from Central Asia to eastern Europe for *C. orientalis* and mountains of northwest Greece and Albania for *C. grandiflora*), *C. bursa-pastoris* has an almost worldwide distribution. The rapid expansion of the shepherd’s purse did not prevent the emergence of three distinct genetic clusters: Asia (ASI), Middle East and northern Africa (ME), and Europe and the Russian Far East (EUR) (Cornille *et al*., 2016). Demographic inferences showed that these three clusters are the result from a range expansion that started either from the Middle East or Europe (the starting point is not known with certainty), and was followed by a consecutive colonization event towards Asia. This recent worldwide spread was, in some cases, likely associated to human migrations, as for instance the spread to eastern Siberia of western European accessions (Cornille *et al*., 2016), or of southern European and Middle Eastern accessions to North America (Hurka & Neuffer, 1997; Cornille *et al*., 2016).

Given conflicting predictions, it remains challenging to explain the apparent ecological success of species, such as *C. bursa-pastoris*, that recently expanded their range. Moreover, adopting a “reciprocal transplant” framework to address this question would be hard to implement as it would require numerous transfers to capture the genetic diversity and the environmental range of *C. bursa-pastoris*. Instead, in order to investigate the joint role of phenotypic plasticity and adaptation in the colonization success of *C. bursa-pastoris*, we implemented an experimental design with three large common gardens located in three contrasted environments. Firstly, in order to reflect the diversity of environmental conditions that *C. bursa-pastoris* faced during its range expansion, we installed two common gardens at extreme latitudes in Eurasia, one in East Asia and one in Northern Europe, and a third one in North America, which lies outside of the native range of the species. Secondly, to capture the demographic history of *C. bursa-pastoris*, we used a comprehensive sampling of populations from Europe, Asia, North Africa, the Middle East and North America. And thirdly, to be able to characterize phenotypic variation and accession’s performances, we measured several life-history and phenological traits, some of which are main fitness components. This provided us with a solid frame-work to: (i) measure the responsiveness of individual accessions to different environments (i.e., assess their phenotypic plasticity) by comparing the performance of each accession across the three common gardens; (ii) estimate the genetic cluster effect associated to past demographic history by comparing the performances of the three genetic clusters across the common gardens; and (iii) assess the variation in responsiveness among accessions of a same cluster using Finlay-Wilkinson regressions (Finlay & Wilkinson, 1963). Evidence of the role of adaptation in the colonization process would be provided by analyzing all the results together and notably by estimating the genotype (genetic cluster) by environment (common garden) interactions. We will discuss these results in the light of *C. bursa-pastoris* colonization history and its accumulated expansion load.

## Materials and Methods

### Common gardens

#### Localization

One common garden was located at Uppsala (59°51’N, 17°38’E, Sweden), the second one at Guangzhou (23°08’N, 113°16’E, China), and the third one at Toronto (43°39’N, 79°23’W, Canada). To account for the sensitivity of *C. bursa-pastoris* to extreme environmental conditions and to guarantee intermediate temperatures, experiments were started at different periods of the year: at Uppsala, the experiment started in early May 2014 for 139 days; at Guangzhou, in early November 2014 for 193 days; and at Toronto, in early June 2014 for 118 days.

#### Environmental conditions

At Uppsala and Guangzhou, environmental conditions were monitored daily at ground level using temperature and humidity sensors (TGP-4017^®^, Tinytag™). At Toronto, temperature and humidity were extracted from a public database (https://www.timeanddate.com). For each location, day length was obtained with the R package *geosphere* (function *daylength*; Hijmans, 2019).

### Plant materials

#### Sampling

We used a collection of 267 accessions from 65 sites distributed across Europe, Asia, North Africa, the Middle East and North America (for more details, see Cornille *et al*., 2016, Kryvokhyzha *et al*., 2016, Kryvokhyzha *et al*. 2019b). In Toronto, due to limited space, we had to subsample this collection down to 160 accessions, and were chosen to share less than 97% of identity-by-state (for further details, see Salcedo, 2015), to maximize the genetic diversity.

#### Genotypes

Genotyping-by-sequencing (GBS) data for the 267 accessions acquired by Cornille *et al*., (2016, https://datadryad.org/stash/dataset/doi:10.5061/dryad.71f99) were used to assign accessions to different clusters (Asia, Middle East or Europe). First, we ran a Multi-Dimensional Scaling analysis (function c*mdscale* from the base R package *stats*) on the GBS data that gave the five first principal components on which we ran an unsupervised k-means algorithm with *K = 3* (function *kmeans* from the base R package *stats*). We then tested its robustness by using a random forest approach (function *randomForest* from the R package *randomForest*; Liaw & Wiener, 2002), with a bootstrap (1000 iterations) of 20% of the dataset (80% being used as a training set). Only accessions that were assigned more than 95% of the iterations to the same cluster were kept for further analyses (Table S1 and Figure S1).

#### Seeds preparation

Before the establishment of the common garden experiment, and to limit maternal effects, seeds collected from the field were first sown and grown under controlled conditions in growth chambers (55% moisture, 22°C, 12h:12h light:darkness cycles). Their offspring (seeds) were then used to establish the common gardens. About 20 seeds per accession were surface-sterilized the same day and germinated in Petri dishes, with MS medium and agar (see protocol in Kryvokhyzha *et al*., 2016). Petri dishes were then stratified for seven days at 4°C in the dark to promote germination. After this cold treatment, Petri dishes were placed in a greenhouse with no additional light or heating, in order to protect seeds from rainfall and to facilitate acclimation to outdoor conditions. Petri dishes were randomized over tables and moved every day to avoid micro-environmental effects, and they were left in the greenhouse until seedlings reached a four-leaf stage.

#### Transplantation

At Uppsala and Guangzhou, once a seedling reached the four-leaf stage, it was thinned to one per pot (7 cm x 7 cm) containing mud (basic soil mixed with water). The pots were left seven days in the greenhouse (watered automatically twice a week) and then they were pierced on the bottom and placed outside in the common garden. The pots were dispatched into six blocks (1m x 3.2m, grids of 9 x 30) arranged at 2 m spacing and containing basic soil. Each block contained exactly one replicate per accession, so that we ended up with six replicates per accession in total. An accession was included in the common garden experiment even if less than six seedlings germinated. In such case, and to keep the same individual density in each block, seedlings from another accession from the same sampling site were planted. At Toronto, after reaching the four-leaf stage, seedlings were directly transplanted from Petri dishes to planting beds (13 cm x 33 cm) containing Promix soil (thoroughly wet). These beds contained exactly one replicate per accession, and we used the same workaround when germination failed. The beds were covered with shade cloth for two days after planting. At each location, apart from the initial shading and watering, we did not provide any support to the seedlings. The experiment lasted until the senescence of the last plant, *i*.*e*., when the last plant dried up but had not yet shed its fruits.

#### Subdividing datasets

We considered three approaches for studying the 267 accessions: for exploratory analyses, we kept the “whole dataset”, accepting potential unbalance; this dataset was thereafter referred to as the whole dataset. Since the interaction effect between common garden and genetic cluster was significant (see Supplementary materials), following the recommendation of Crawley (2012), we chose to analyze each common garden separately; this dataset was thereafter referred to as the “common garden dataset”. Finally, in order to mimic a reciprocal transplant experiment and test for signature of local adaptation, we considered a sub-dataset made only of accessions from Sweden and South China (12 accessions in total, sampled close to Uppsala or Guangzhou), and excluding their phenotype measurement in Toronto; this dataset was thereafter referred to as the “local adaptation dataset”.

### Statistical analyses

#### Phenotypes

After senescence, we recorded for each plant the *height of the highest inflorescence* (in dm), the *number of basal inflorescences*, and the *number of fruits along a section of 1 dm* in the middle of the main inflorescence. In order to compare the overall performances, we considered the total number of fruits per individual as a proxy of fitness (denoted *w*), that were measured differently in the common gardens: at Uppsala and Guangzhou, we considered a composite index of fitness, computed as the product of the aforementioned three traits (*number of basal inflorescences, height* and *number of fruits along a section of 1 dm*). At Toronto, the total number of fruits was directly measured for each individual.

Three phenological traits were also measured, *bolting time* (*i*.*e*., time until differentiation of the bud from vegetative parts indicating the initiation of the reproductive period), *flowering time* (*i*.*e*., time until the appearance of the first opened flower) and *senescence time* (*i*.*e*., time until the drying state of the plant). In addition, *flowering time span* was computed as being the inter-event time between *flowering time* and *senescence time*. At flowering time, we also recorded the rosette leaf number and the maximum diameter of the rosette. Finally, at Uppsala and Guangzhou, germination time was also measured.

#### Statistical modeling

For the statistical analyses, we fitted the following generalized linear mixed model to the data:

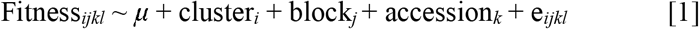

where *μ* is the overall mean, ‘cluster’ is the fixed effects of the genetic cluster, ‘block’ and ‘accession’ are uncorrelated random effects (‘accession’ being nested into ‘cluster’), and ‘e’ is the residual. We used a log link function and the residual distribution was fitted to a Negative Binomial distribution (function *glmer*.*nb* of R package *lme4*; see Bates *et al*., 2015). This model was thereafter referred to as the “GLMM model”. For phenological traits, we fitted the GLMM model to the four inter-event durations (between sowing, germination, bolting, flowering, and senescence), square-rooted, with the starting date of each time span as an additional explaining variable, and with the residual fitted to a normal distribution (function *lmer* of R package *lme4*). Statistical significance of the cluster effect was assessed with a type II Wald chi-square test, and the difference between genetic clusters was assessed with Tukey’s HSD test.

#### Finlay-Wilkinson regression (FWR)

We further investigated the specificity of each genetic cluster by performing a Finlay-Wilkinson, 1963), enabling us to characterize individual genotype responses for a trait across different environments. Following the formulation of Lynch & Walsh (1998), a FWR for one accession can be expressed as follows:

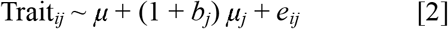

where ‘Trait’ is the focal trait (fitness or phenological traits), *i* is the block, *j* is the common garden, *μ*_*j*_ is the average value of the trait at the common garden *j, b*_*j*_ is the regression factor of *μ*_*j*_ and *e*_*ij*_ is the residual which is assumed to be normally distributed. We performed as many regressions as the number of accessions. This model was thereafter referred to as the FWR model.

The intercept (*μ*) can be interpreted as the accession’s performance in a poor environment, and the slope (*b*_*j*_) can be viewed as the accession’s relative responsiveness to environmental changes (the steeper the slope, the more responsive the accession; Lynch & Walsh, 1998). By definition, across the whole dataset, we expect a null average intercept (*μ = 0*) and a unit average slope (*b*_*j*_ *= 0*). Each genetic cluster can, however, have its own specificity and deviate from the global trend. We used FWR to assess whether a genetic cluster was homogeneous, *i*.*e*., with a trend followed by every individual belonging to this cluster, or heterogeneous and actually driven by some “outliers”. For intercept (*μ*) and slope (*b*), the statistical significance of difference in average or variance between genetic clusters was assessed with a pairwise Welch’s *t*-test or a Fisher’s *F*-test, respectively.

## Results

Statistical interaction between “common garden” and “genetic cluster” was significant for most traits (Table 1 and Supplementary materials), therefore we focused our analyses on the common garden dataset with the GLMM model; the use of the whole dataset was restricted to the description of phenotypic variation (Table 1 and Figures 1 to 3). Due to experimental issues (*e*.*g*., failed germination, climatic events), common garden experiments ended up with an unbalanced block design: the whole dataset was left as is, but we shrank the common garden dataset to balance it, ending up with 186 accessions in Uppsala, 86 in Guangzhou and 116 in Toronto (Table S1). Also, due to these issues, the number of accessions common to all three common gardens in the common garden dataset was reduced to a total of 34 accessions (12 Asian, 9 European and 13 Middle Eastern): we therefore fitted the FWR model on the (unbalanced) whole dataset filtered for accessions for which a fitness score was available in all three common gardens, ending up with a total of 114 accessions (46 Asian, 46 European and 22 Middle Eastern).

**Table 1.** Analysis of variance of model [S1] (see Supplementary materials). Statistics (*χ*^2^), degree of freedom (df) and p-values of the type II Wald chi-square test. Only variables that were measured in all the common gardens are kept. FP: time between bolting and flowering; SP: time between flowering and senescence. The distributions of the residual are normal distributions. Significance levels are : *p\*\*\** < 0.001; *p** <* 0.01; *p\** < 0.05; *p*^*n*.*s*.^ > 0.05. Since germination time was not monitored in Toronto, analysis of variance was not assessed for germination time and the inter event between bolting time and germination time.

|  | Genetic cluster |  | Common garden |  | Genetic cluster x common garden |  |
| --- | --- | --- | --- | --- | --- | --- |
| | $\chi^2$ | df | $\chi^2$ | df | $\chi^2$ | df |
| Fitness (Nb. fruits) | 67.1*** | 2 | 128.5*** | 2 | 135.1*** | 4 |
| Height | 109.9*** | 2 | 248.9*** | 2 | 122.0*** | 4 |
| Nb. primary branches | 3.2<sup>n.s.</sup> | 2 | 420.8*** | 2 | 194.7*** | 4 |
| Nb. rosette leaves | 227.5*** | 2 | 358.7*** | 2 | 517.5*** | 4 |
| Rosette diameter | 88.8*** | 2 | 309.4*** | 2 | 45.4*** | 4 |
| FP | 66.2*** | 2 | 656.8*** | 2 | 2.2<sup>n.s.</sup> | 4 |
| SP | 10.4*** | 2 | 1445.2*** | 2 | 6.1<sup>n.s.</sup> | 4 |

**Figure 1.**
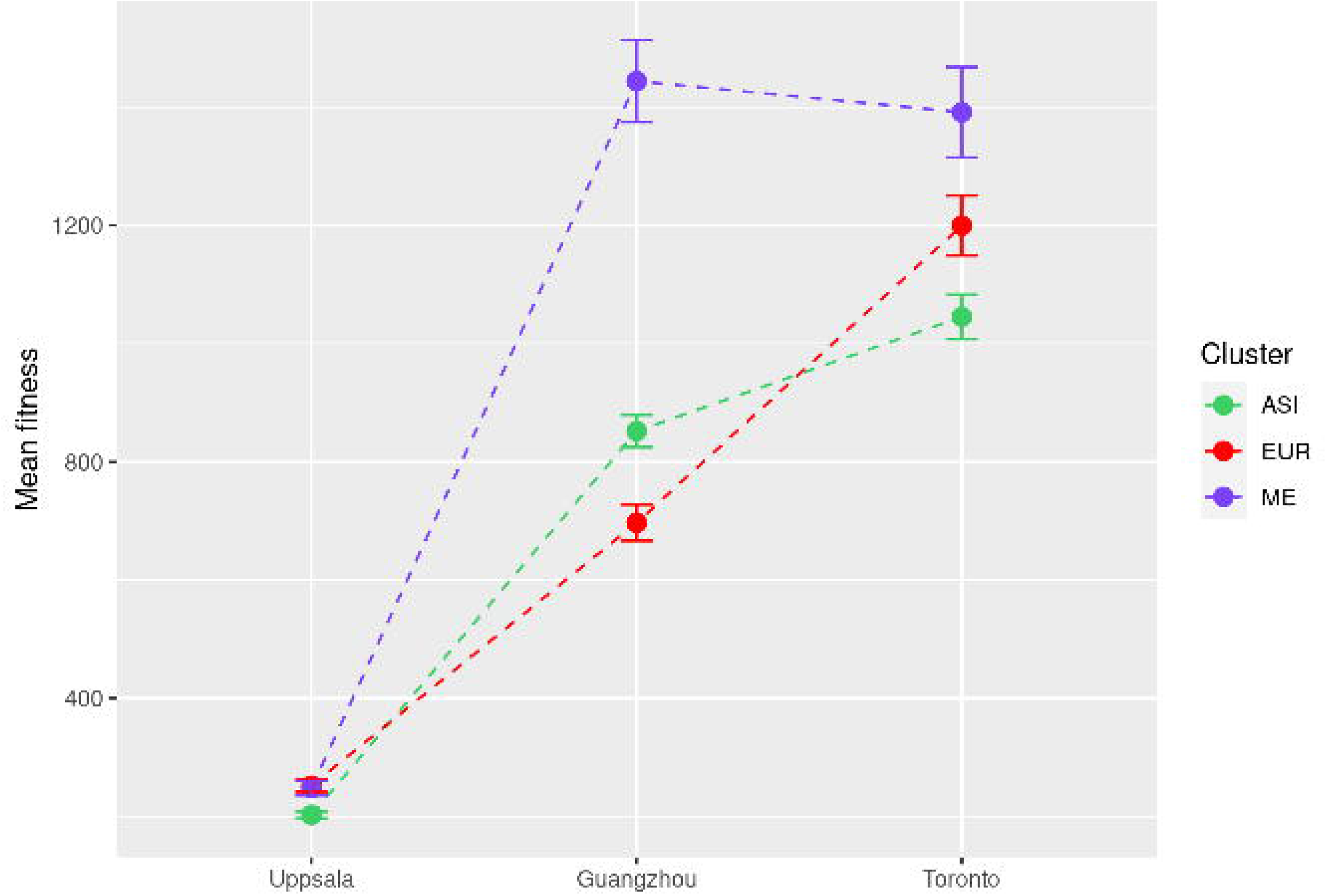
Interaction plot of fitness in *C. bursa-pastoris* between the three common gardens (Uppsala, Guangzhou and Toronto, on the x-axis) and the genetic clusters (ASI in green, EUR in red, and ME in blue), using the whole dataset. Error bars correspond to standard errors.

At the time of the experiment (2014), environmental conditions (in terms of temperature, humidity and day length) strongly differed among the common gardens (Figure 1). In Uppsala, accessions grew in rather cold and wet climatic conditions (15.2 ± 5.1°C, 76.8 ± 14.1%); in Toronto they grew under a relatively warm and dry climate (20.7 ± 3.1°C, 71.2 ± 8.9%); finally, in Guangzhou they grew in a warm and humid weather (19.3 ± 5.0°C, 81.1 ± 11.4%). The photoperiod and the magnitude of its variation during the experiment also strongly differed among common gardens, day length being the longest in Uppsala (16.9h ± 1.7h, ranging from 13h to 18.8h), the shortest in Guangzhou (11.9h ± 1.0h, ranging from 10.7h to 13.5h), and intermediate in Toronto (14.3 ± 1.1h, ranging from 12h to 15.4h). For further details, see Tables S2 to S4 and Figures S2 to S5.

### Environmental variation as the main explanatory factor of the phenotypic variation of *Capsella bursa-pastoris*

Most of the phenotypic variation was explained by the common garden, with a strong environmental effect (Figure 1, Table 1).

Accessions showed contrasted phenotypes among common gardens (Figures 2 and S6), highlighting their phenotypic plasticity. In Toronto, accessions tended to be more “bushy” with relatively more branches and more rosette leaves, whereas in Guangzhou, accessions tended to be relatively taller. In Uppsala, accessions were globally smaller and performed poorly for most phenotypic traits. The common garden effect was even more pronounced on phenology than on life history traits (*χ*^*2*^_*pheno*_ *=* 1051 ± 557.5; *χ*^*2*^_*life history traits*_ = 293.3 ± 111.7; all *p* < 0.001; Table 1). In particular, *bolting time, flowering time* and *senescence time* were longer in Guangzhou (respectively 61.3 ± 13 days, 71.5 ± 13.1 days, 138 ± 9.5 days) than in Toronto (28.0 ± 4.8 days, 31.9 ± 7.5 days, 79.8 ± 9.5 days) and Uppsala (30.9 ± 6.1 days, 36.4 ± 5.6 days, 85.3 ± 7.6 days).

**Figure 2.**
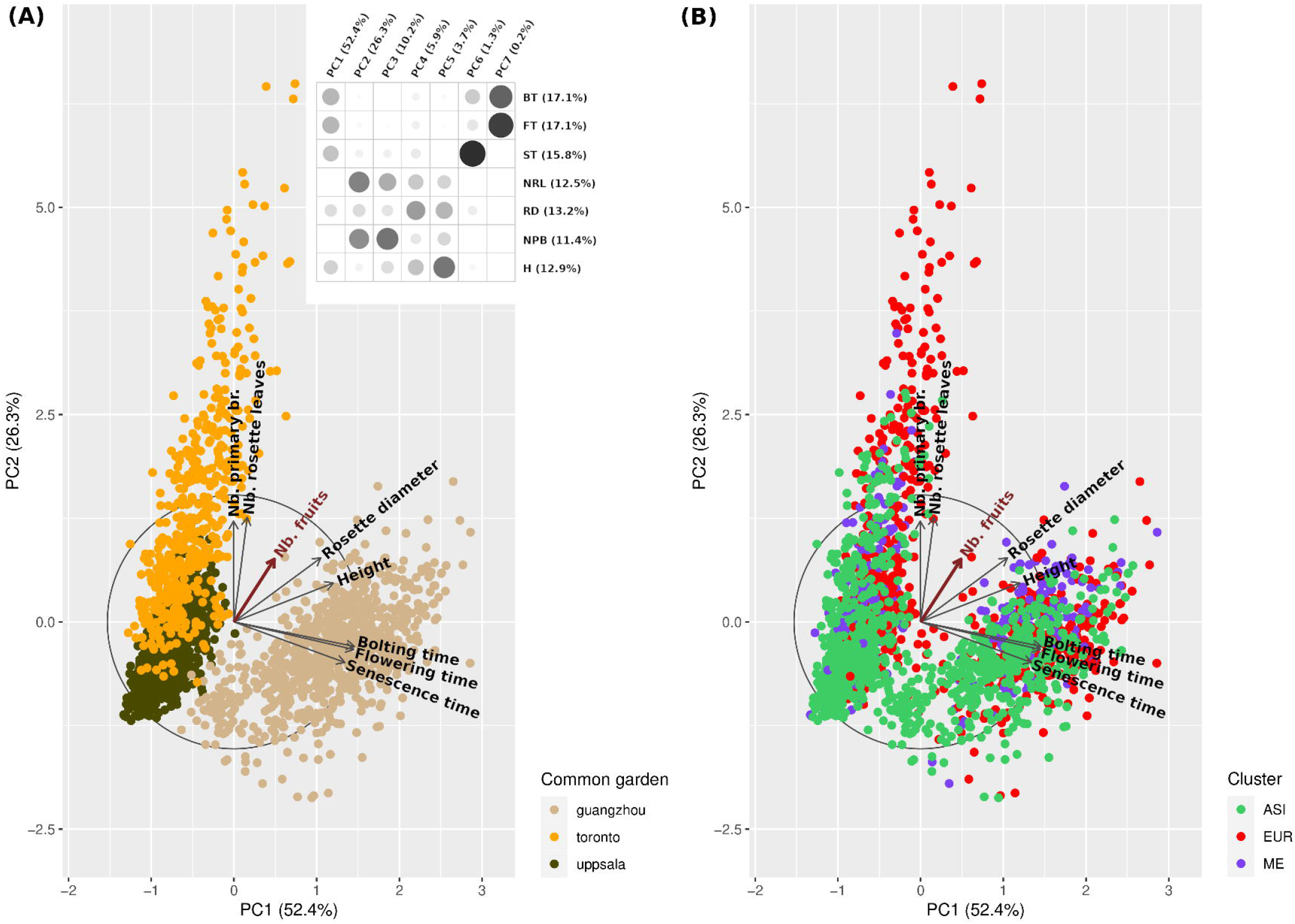
Principal Component Analysis of the phenotypes of accessions of *C. bursa-pastoris*: bolting time (BT), flowering time (FT), senescence time (ST), number of rosette leaves (NRL), rosette diameter (RD), height of the main inflorescence (H), and the number of primary branches (NPB), using the whole dataset (267 accessions). The number of fruits (the fitness) was added as a supplementary variable. PC1 captured 52.4% of the variance, and PC2 captured 26.3%. The box on the top right corner of (**a**) highlights the correlation of each variable to the corresponding axis: the darker and bigger the circle, the higher the correlation (in brackets, the percentage of contribution of each variable to the first two PCs). Both (**a**) and (**b**) are the same PCA, only the color label changes. (**a**) PCA labeled according to the common gardens (Uppsala in brown, Toronto in orange, and Guangzhou in beige): common gardens appeared to be clearly delimiting three groups. (**b**) PCA labeled according to the genetic clusters (ASI in green, EUR in red, and ME in blue): genetic clusters did not appear to be clearly separating the dataset.

**Figure 3.**
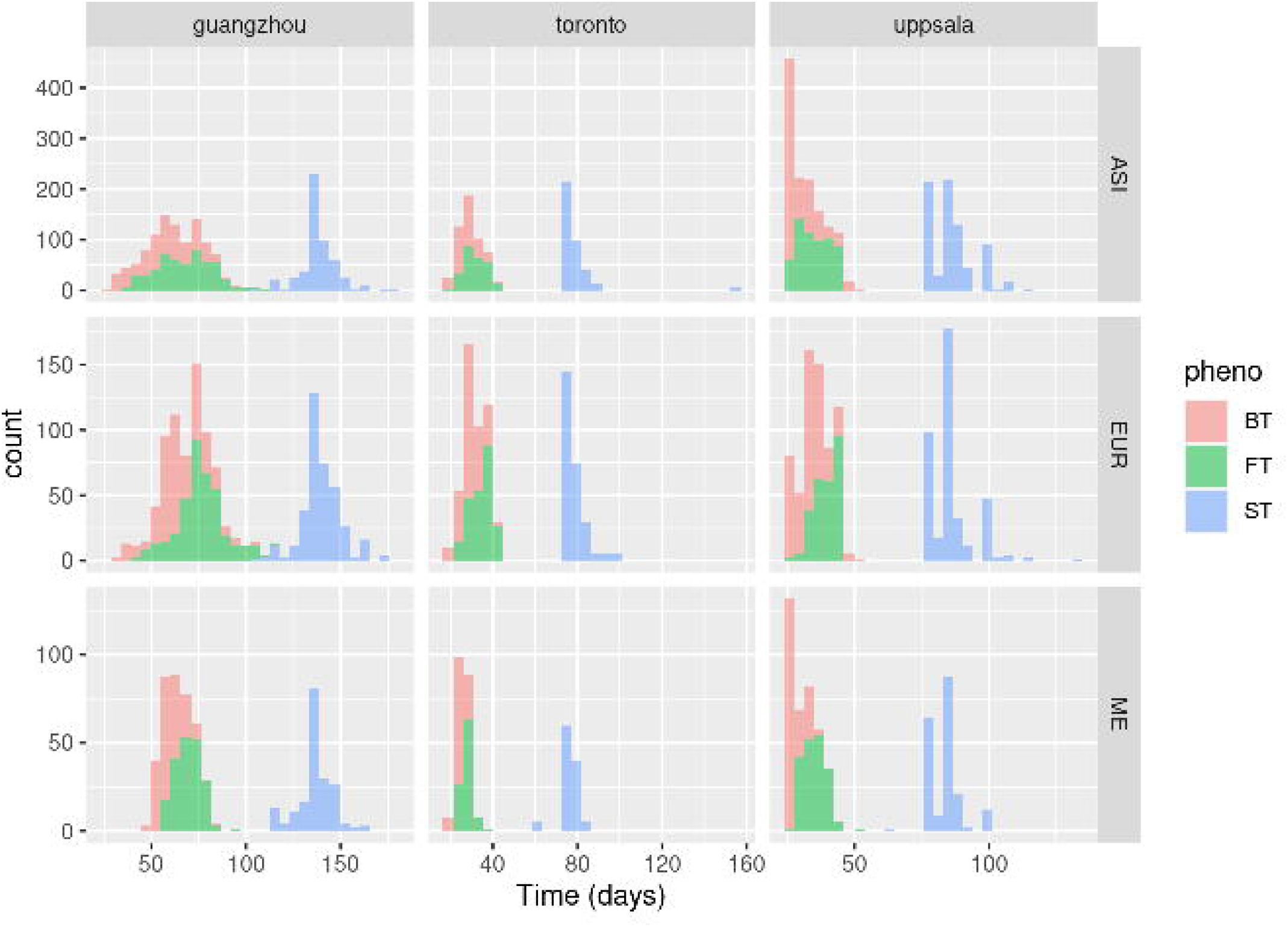
Histograms of the phenological traits of *C. bursa-pastoris* (bolting time as BT, flowering time as FT, and senescence time as ST) according to the common garden and genetic cluster, using the whole dataset. Germination time was not plotted here for consistency with Toronto.

**Figure 4.**
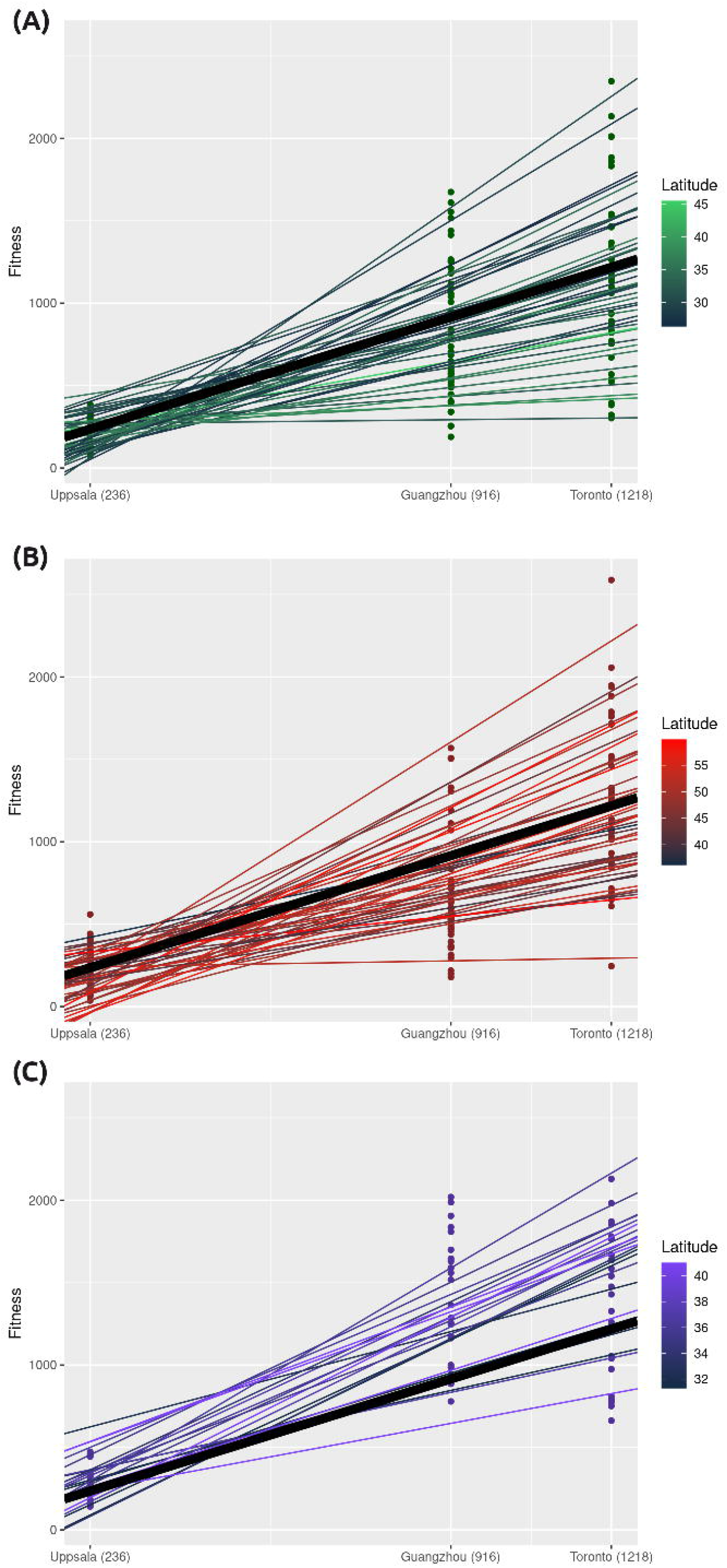
Finlay-Wilkinson regressions of fitness (the FWR model) of *C. bursa-pastoris*. On the x-axis, the mean population fitness at each common garden; on the y-axis the mean fitness of each accession. Each dot is an accession at a specific common garden, and each line is a FWR for one accession. The bold black line is the population average FWR. (A) FWR of accessions belonging to the cluster ASI (green); (B) FWR of accessions belonging to the cluster EUR (red); (C) FWR of accessions belonging to the cluster ME (blue).

More importantly, environmental effect on fitness was significant (*χ*^*2*^ = 128.5, df = 2, *p* < 0.001, type II Wald chi-square test; Table 1): fitness was much lower on average in Uppsala (*w* = 224 ± 178) than in Guangzhou (*w* = 918 ± 753) and Toronto (*w* = 1161 ± 788). Toronto presented the most favorable conditions in terms of temperature and day length, two factors *C. bursa-pastoris* is known to be particularly sensitive to.

### Genetic effect associated to the past demographic history significantly explained variation within a common garden

Genetic effect comes second and explains the phenotypic variation within each common garden (Figures 1 and 2).

#### Phenotypes

Within each common garden, except for the number of primary branches, phenotypic variation showed a strong genetic effect (Table 2). The cluster effect for the number of primary branches was not significant in Uppsala, and strongly significant (*p* < 0.01) in Guangzhou and Toronto. Surprisingly, the cluster effect for rosette characteristics (number of leaves and diameter) was much more significant than that for height of the main inflorescence or number of primary branches (Table 2). On average, the size and the number of leaves of the European accessions were much higher (respectively 188.6 ± 55.5 mm and 40.8 ± 30.4) than those of the Middle Eastern accessions (respectively 172 ± 54.8 mm and 19.3 ± 8.4) and of the Asian accessions (121.0 ± 56.5 mm and 19.5 ± 11.7).

**Table 2.**
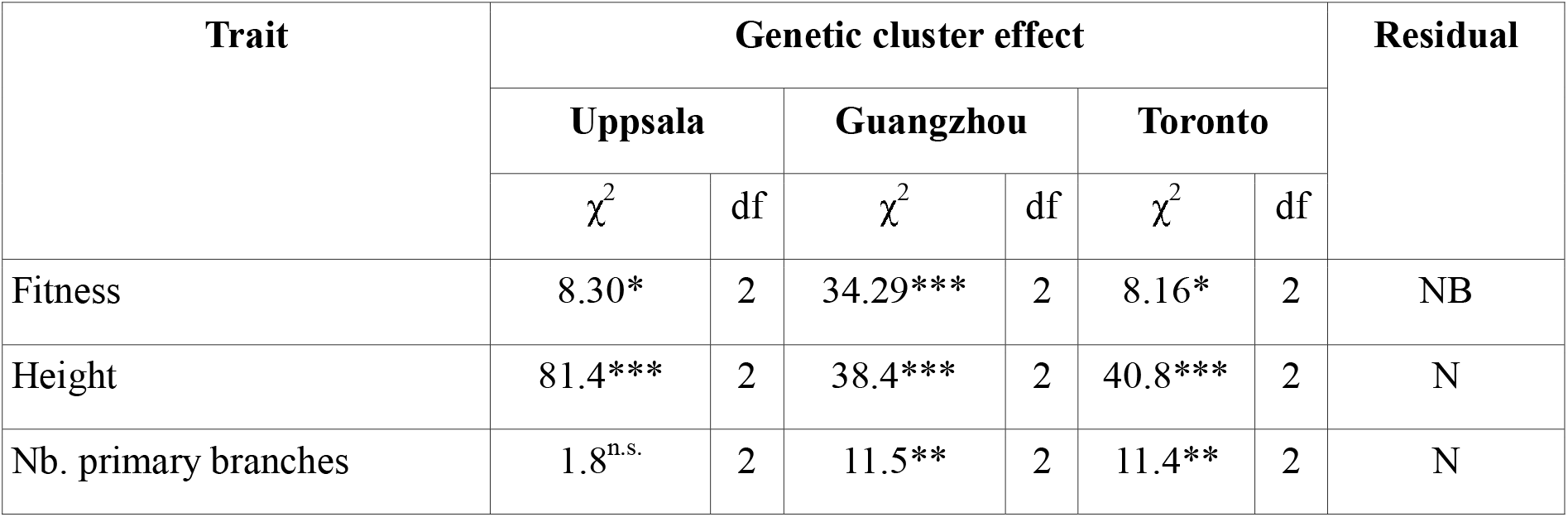
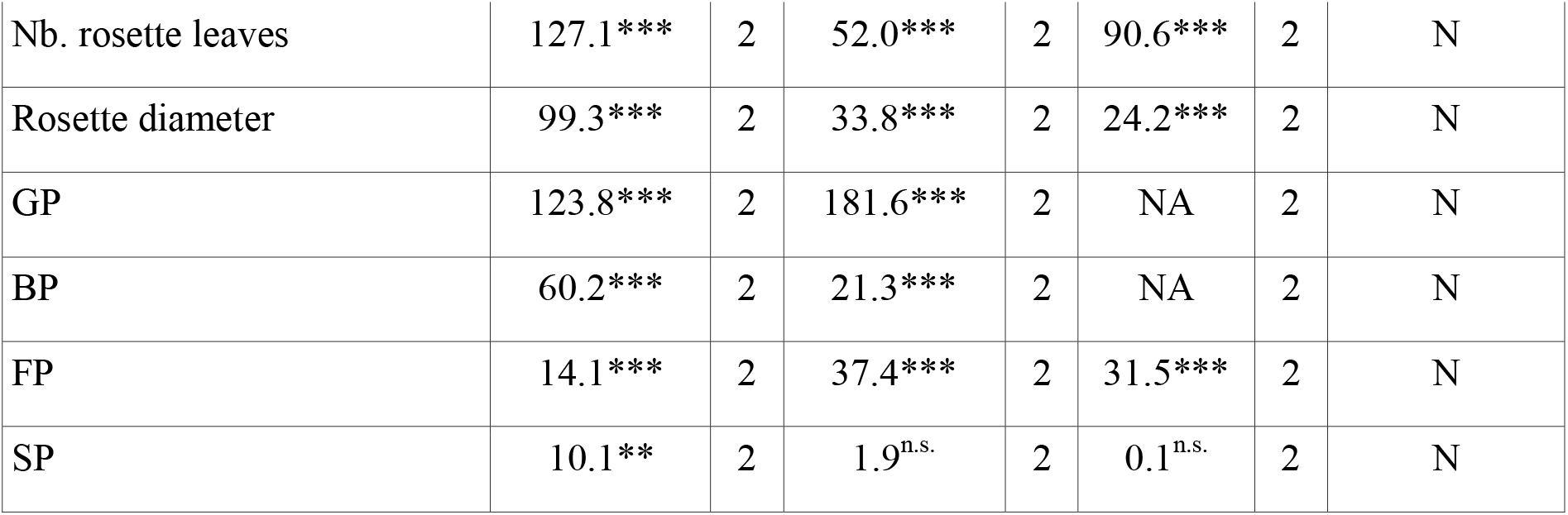
Analysis of variance of the GLMM model. Statistics (*χ*^2^), degree of freedom (df) and p-values of the type II Wald chi-square test. GP: time between sowing and germination; BP: time between germination and bolting; FP: time between bolting and flowering; SP: time between flowering and senescence. The distributions of the residual are : negative binomial (NB), or Normal (N). Significance levels are : *p\*\*\** < 0.001; *p** <* 0.01; *p\** < 0.05; *p*^*n*.*s*.^ > 0.05. Since germination time was not monitored in Toronto, analysis of variance was not assessed in Toronto for GP and BP.

| Trait | Genetic cluster effect |  |  |  |  |  | Residual |
| --- | --- | --- | --- | --- | --- | --- | --- |
|  | Uppsala |  | Guangzhou |  | Toronto |  |  |
| | $\chi^2$ | df | $\chi^2$ | df | $\chi^2$ | df | |
| Fitness | 8.30* | 2 | 34.29*** | 2 | 8.16* | 2 | NB |
| Height | 81.4*** | 2 | 38.4*** | 2 | 40.8*** | 2 | N |
| Nb. primary branches | 1.8<sup>n.s.</sup> | 2 | 11.5** | 2 | 11.4** | 2 | N |
| Nb. rosette leaves | 127.1*** | 2 | 52.0*** | 2 | 90.6*** | 2 | N |
| Rosette diameter | 99.3*** | 2 | 33.8*** | 2 | 24.2*** | 2 | N |
| GP | 123.8*** | 2 | 181.6*** | 2 | NA | 2 | N |
| BP | 60.2*** | 2 | 21.3*** | 2 | NA | 2 | N |
| FP | 14.1*** | 2 | 37.4*** | 2 | 31.5*** | 2 | N |
| SP | 10.1** | 2 | 1.9<sup>n.s.</sup> | 2 | 0.1<sup>n.s.</sup> | 2 | N |

#### Phenology

Phenological variation also showed a strong and significant cluster effect (Table 2). The Asian cluster exhibited a late germination, early bolting, early flowering and late senescence, therefore a long flowering time span. In contrast, the European cluster germinated early, but bolted, flowered and withered late (Table S5). European accessions’ late flowering partly explains their larger rosette, since they had more time for vegetative growth. Accessions from the Middle-East genetic cluster did not follow a global trend (Table S5), deploying again a certain amount of plasticity in their phenological response.

For *bolting time*, mean intercept in the FWR model was significantly lower for the Middle Eastern than for the Asian and the European clusters (all *p* < 0.05, Welch’s *t*-test; Table S6): in an environment pushing *C. bursa-pastoris* to bolt early on average, ME bolted even earlier.

#### Fitness

Variation in fitness also showed a strong and significant genetic effect (Figure 1 and Table 2). The Middle Eastern cluster globally outperformed the other clusters, with an overall mean performance of *w* = 946 ± 916, compared to the Asian (*w* = 565 ± 613) and the European (*w* = 652 ± 685) genetic clusters. At each common garden, its performance is either significantly higher than another cluster, or not significantly smaller than another one (Table 3). The Middle Eastern cluster deployed a significantly higher phenotypic variance during this experiment than the other clusters (all *p* < 0.001, Fisher’s F-test; Table S7), and also showed a significantly higher responsiveness (slope in the FWR model; all *p* < 0.01, Welch’s t-test; Table 4), corroborating the hypothesis of a higher plasticity of the Middle Eastern cluster.

**Table 3.** Estimate of genetic cluster effect in the GLMM model, in log-scale at each common garden (Uppsala, Guangzhou, Toronto). Estimates of EUR and ME effects are relative to ASI effect; Tukey contrast analysis of fitness in the GLMM model: statistics and p-values of the Tukey’s HSD test. Significance levels are : *p\*\*\** < 0.001; *p** <* 0.01; *p\** < 0.05; *p*^*n*.*s*.^ > 0.05.

| Cluster | Uppsala |  |  | Guangzhou |  |  | Toronto |  |  |
| --- | --- | --- | --- | --- | --- | --- | --- | --- | --- |
|  | Estimate | Tukey's HSD |  | Estimate | Tukey's HSD |  | Estimate | Tukey's HSD |  |
|  |  | EUR | ME |  | EUR | ME |  | EUR | ME |
| ASI | 4.56 | -2.6* | -1.8<sup>n.s.</sup> | 6.12 | 1.8<sup>n.s.</sup> | -4.5*** | 6.13 | -1.4<sup>n.s.</sup> | -2.8* |
| EUR | +0.21 | - | 0.3<sup>n.s.</sup> | -0.20 | - | -5.6*** | +0.14 | - | -1.8<sup>n.s.</sup> |
| ME | +0.18 | - | - | +0.50 | - | - | +0.37 | - | - |

**Table 4.** Mean and standard deviation of responsiveness (slope) and performance in poor environment (intercept) of fitness (the FWR model), for each genetic cluster (ASI, EUR, and ME); pairwise Welch’s *t*-test between mean values of genetic clusters: statistics (T), degree of freedom (Df) and p-values. Significance levels are : *p\*\*\** < 0.001; *p** <* 0.01; *p\** < 0.05;*p*^*n*.*s*.^ > 0.05.

| Cluster | Intercept |  |  |  |  | Slope |  |  |  |  |
| --- | --- | --- | --- | --- | --- | --- | --- | --- | --- | --- |
|  | Mean ± SD | Welch's *t*-test |  |  |  | Mean + SD | Welch's *t*-test |  |  |  |
|  |  | EUR |  | ME |  |  | EUR |  | ME |  |
|  |  | T | Df | T | Df |  | T | Df | T | Df |
| ASI | 6.70 ± 183.5 | 0.94<sup>n.s.</sup> | 90.0 | -0.38<sup>n.s.</sup> | 40.4 | 0.91 ± 0.47 | -0.5<sup>n.s.</sup> | 90.0 | -3.5** | 53.2 |
| EUR | -29.92 ± 189.0 | - | - | -1.1<sup>n.s.</sup> | 41.5 | 0.96 ± 0.44 | - | - | -3.1** | 50.4 |
| ME | 25.47 ± 188.6 | - | - | - | - | 1.27 ± 0.36 | - | - | - | - |

### The Middle Eastern accessions had a more homogeneous response to environmental variation than Asian or European accessions

#### Phenology

The Middle-Eastern cluster showed, according to the FWR model, a lower variance in their phenological behavior: the variance of the slope for *flowering time, flowering time span* and *bolting time* was significantly smaller than that of the Asian and the European accessions (all *p* < 0.001, Fisher F-test; Table S6), therefore suggesting that the Asian and European clusters are far more diverse than the Middle Eastern one.

#### Fitness

Variance of the intercept in the FWR model was too large compared to its mean value to be interpreted, given the coefficients of variation of 27.4 for the Asian, 6.3 for the European and 7.7 for the Middle Eastern (Table 4) clusters. However, the slope was more discriminant: the Middle Eastern cluster had a significantly higher (all *p <* 0.01, Welch’s *t*-test; Table 4) responsiveness (1.27 ± 0.36) than the Asian (0.91 ± 0.47) and the European (0.96 ± 0.44) clusters. Slope variance of the Middle-Eastern cluster was smaller than for the two other clusters,, although not significantly (Table S6).

#### Linking fitness and phenology

Surprisingly, Pearson’s correlation between different FWRs was weak (absolute values < 0.2 if putting ME aside; Table 5), suggesting that phenology explained only a small fraction of the variance of fitness. However, for the Middle-Eastern cluster, Pearson’s correlation between *fitness* and *flowering time* or between *fitness* and *flowering time span* was equal to *r* = 0.30 and *r* = 0.50 respectively, suggesting that accessions that were plastic for part of the phenology were likely also plastic for the number of fruits.

**Table 5.** Correlation between different Finlay-Wilkinson regressions (slope and intercept) on fitness and flowering time (FT), or flowering duration (SP).

| Cluster | Intercept |  | Slope |  |
| --- | --- | --- | --- | --- |
|  | FT | SP | FT | SP |
| ASI | 0.15 | 0.03 | 0.06 | -0.07 |
| EUR | -0.10 | 0.09 | -0.19 | 0.10 |
| ME | -0.15 | 0.14 | -0.50 | 0.30 |

### Hints of signature of local adaptation when mimicking a reciprocal transplant

In the presence of local adaptation, we expect a higher fitness of accessions in their own geographical range. In our case, we expected a higher relative fitness of the European accessions in Uppsala, and a higher relative fitness of the Asian accessions in Guangzhou.

#### With the common garden dataset

Whereas the Middle Eastern cluster globally outperformed the other clusters, the European cluster (*w* = 252 ± 216) significantly outperformed the Asian cluster (*w* = 203 ± 153) in Uppsala (*z* = 2.65, *p <* 0.05, Tukey’s HSD test; Table 3). However, although a trend of the Asian cluster (*w* = 852 ± 634) outperforming the European cluster (*w* = 697 ± 604) in Guangzhou can be seen, this difference was not significant (*z* = -1.83, *p* = 0.16, Tukey’s HSD test; Table 3).

#### With the local adaptation dataset

The pattern of local adaptation was actually even less clear when considering the local adaptation dataset: the performances of the South Chinese and Swedish accessions did not differ significantly. Indeed, in Guangzhou, though the performance of Chinese accessions (*w* = 926 ± 598) tended to be higher than that of Swedish accessions (*w* = 524 ± 533), the difference was not significant (*z* = -1.379, *p* = 0.17, Tukey’s HSD test); likewise, in Uppsala, difference in terms of performance between Chinese accessions (*w* = 208 ± 130) and Swedish accessions (*w* = 210 ± 193) was not significant (*z* = -0.752, *p* = 0.45, Tukey’s HSD test). One accession from Sweden outperformed all Chinese accessions though, suggesting that the lack of a significant signature of local adaptation might be caused by the limited number of accessions sampled (N_EUR_ = 3 and N_ASI_ = 9).

## Discussion

Overall, the location of the common gardens explained a large part of the phenotypic variation of *C. bursa-pastoris*, suggesting that the underlying mechanism of its success is probably a high degree of phenotypic plasticity. To a lesser extent, genetic clusters explained phenotypic variance within common gardens. Assuming that populations in the colonization front gradually undergo a shift of their prior phenotypic plasticity to adaptation to local conditions, *C. bursa-pastoris* seems to be in this very transient state. As a matter of fact, the hypothesis of the presence of a transient state in invasive species could explain most of the discrepancies in the literature about the prevalence of either phenotypic plasticity or adaptation. Indeed, if the colonization event is recent, it is possible, like in our study or in VanWallendael *et al*. (2018), that signatures of local adaptation are not yet observable (but see Godoy *et al*., 2011; Ducatez *et al*., 2016).

### The overall success of the Middle Eastern accessions might obscure the scale of local adaptation

*C. bursa-pastoris* is a self-fertilizing species that underwent a relatively recent worldwide expansion, and has a low genetic variation that could, by itself, explain the lack of local adaptation signature in the present study. *C. bursa-pastoris* and its diploid relatives are eminently ruderal and even a casual observation of their habitats highlights the importance of very local factors for their establishment, such as soil disturbance (Orsucci *et al*., *in press*). Nonetheless *C. bursa-pastoris* is genetically structured across its worldwide distribution (Cornille *et al*., 2016), and therefore, to account for this structure, we needed to implement a large-scale experiment with a large number of accessions. Thus, perhaps more than in other species, geographical scale is crucial in studies of adaptation in the *Capsella* genus and what is observed at a given scale may not apply at another one, as exemplified by Neuffer (1996).

To the best of our knowledge, however, there has not been a general study of the local ecological niches of the shepherd’s purse. Our own observations and the few studies available, in particular the study by Caullet (2011) that exhaustively studied populations of *C. bursa-pastoris* and *C. rubella* across a 4 km^2^ agricultural landscape of Central France over two years, indicate that *C. bursa-pastoris* is actually restricted to highly disturbed environments (hedges of fields, for instance). Within such environments it does not seem to be confined to specific types of soil though (Aksoy *et al*., 1998; Neuffer *et al*., 2018). The large spectrum of climatic environments covered by the accessions included in our experiment also suggests that insufficient sampling is unlikely to explain the lack of a strong local adaptation pattern.

As shown above, though the effect of either environment or genetic background seems clear, the interaction between genetics and environment is complex. When considering all common gardens and all clusters at once, there is no clear evidence of local adaptation, mainly because the Middle Eastern cluster outperformed the other clusters everywhere. The Middle Eastern cluster is less variable for the traits considered here, and is consistently better and more responsive to variation in environmental conditions than those from the two other clusters. One of the possible explanations of this complex pattern, obscuring a potential signature of local adaptation, is that there is no or few local adaptation yet: an absence of local adaptation coupled with a pronounced plasticity have already been observed in other invasive species at the colonization front, such as in *Reynoutira japonica* (VanWallendael *et al*., 2018) or in clonally reproducing macrophytes (*Egeria densa, Elodea canadensis* and *Lagarosiphon major*: Riis *et al*., 2010).

It is also worth pointing out that the absence of evident GxE effect may be due to the fact that we did not consider the whole life cycle. Recently, Orsucci *et al*. (2020) emphasized that studying only one component of fitness could be insufficient and potentially misleading if interpreted as fitness, since they do not account for possible trade-offs between fitness components. Germination rate of the mother plant was not measured here, and is the only part of the life cycle that was controlled in a growth chamber. Accounting for the germination rate might change the ranking among clusters, as shown in Orsucci *et al*. (2020), we can thus only say that the Middle Eastern cluster outperformed others in terms of vegetative growth and fruit production, but whether it also does in term of fitness remains to be tested.

### A species with a worldwide distribution might present a complex genetic structure and local adaptation pattern

The absence of local adaptation signature could also be explained by the marginality of our common gardens in deference to the species distribution described by Hurka *et al*. (2012). Indeed, Uppsala is close to the northern boundary of the distribution of European accessions; Guangzhou, likewise, is close to the southern boundary of the distribution of Asian accessions; and Toronto is even outside the natural species distribution. Environmental conditions can be extreme in Uppsala and Guangzhou compared to other locations of the same continent. The extreme localization of our common gardens could thus have hidden the importance of local adaptation among clusters, as discussed in Klisz *et al*. (2019). We attempted to mimic a reciprocal transplant experiment by only considering Swedish and South Chinese accessions at Uppsala and Guangzhou. However, the results were mixed, with no apparent signature of local adaptation (i.e., no apparent genetic cluster effect). To rigorously address this problem in further studies, it will be necessary to either (i) increase the amount of actual local accessions, so that the common gardens are less marginal relative to the set of accessions sampled, e.g., more Swedish and South Chinese accessions; or (ii) consider several common gardens in a smaller geographical scale, *e*.*g*., two or more common gardens in Europe.

The absence of evident G x E effect for all of the clusters leads to question about the geographical scale at which local adaptation occurs. Although the populations we considered are at the right genetic scale in the sense that our samples do capture the major genetic clusters in Eurasia (Cornille *et al*., 2016; Kryvokhyzha *et al*., 2019a), local adaptation might occur at a much more restricted geographical scale, as pointed out above. First, *C. bursa-pastoris* is a ruderal species and probably a human commensal too, as suggested by its distribution, and therefore it grows primarily, if not uniquely, in very specific, disturbed environments, that will share common characteristics under different latitudes. Second, *C. bursa-pastoris* is mainly self-fertilizing which implies that gene flow occurs through seeds rather than pollen, thus slowing down its scattering. The hypothesis of adaptation to micro-environment rather than macro-environment is also supported by the fact that phenology is mostly influenced by environmental variation, and only secondly by genetics.

The apparent lack of local adaptation observed in the present study at the phenotypic level can be related to the results obtained by Kryvokhyzha *et al*. (2016). Using RNA-Seq data from 24 accessions originating from the three clusters, Kryvokhyzha and co-authors showed that the genes that were significantly differentially expressed between the three clusters, were no longer so once one had corrected the data for the genetic relationship between the three clusters. The differentiation could then have been mostly neutral or both neutral and adaptive in which case demographic history and adaptive differentiation were highly correlated and correcting for the first one would remove the effect of the latter. The fitness results obtained here would support at least a significant contribution of neutral processes. As in the RNA-Seq experiment, the present data does not rule the presence of significant local adaptation at a more local scale. However, even at that scale local adaptation does not seem to be pronounced in *C. bursa-pastoris* (Caullet, 2011; Huang *et al*., 2018).

It is therefore likely that *C. bursa-pastoris* was able to cope with diverse environments primarily through phenotypic plasticity. Caullet (2011) also found that plasticity best explained fitness across soil niches when they planted both *C. bursa-pastoris* and *C. rubella* in four different disturbed environments where both species are naturally found: along paths, hedges of the paths, or within a transition zone between paths and fields and inside the internal hedge of the fields. The transplant environment did not significantly affect the fitness of *C. bursa-pastoris*. As a whole, as suggested by Caullet (2011), phenotypic plasticity was higher in *C. bursa-pastoris* than in *C. rubella* but there was no evidence of strong local adaptation in *C. bursa-pastoris*. Another possible explanation for the absence of signature of local adaptation is that *C. bursa-pastoris* failed to adapt locally due to its high genetic load. Indeed, *C. bursa-pastoris* is a selfer, with limited genetic diversity and it might have faced a drift barrier and lacked the initial effective population size needed to express local adaptation.

### Phenological traits, between plasticity and adaptation

Controlling the timing of phenology to cope with biotic and abiotic stresses is the response adopted by many plant populations, especially those with an annual life cycle. Adaptation is one of the possible evolutionary responses, as seen in temperate crop species (e.g., wheat, barley or pea) with distinct vernalization strategies in different varieties (Roux *et al*., 2006). As to *C. bursa-pastoris*, in agreement with previous studies (Neuffer & Hurka 1986; Neuffer, 1990), our experiment revealed that its phenological response was mainly through phenotypic plasticity, with a rather equivalent responsiveness among genetic clusters. Although Neuffer & Hurka (1986) evoked a “general purpose genotype” (a term borrowed from Baker, 1965) for germination strategy, we could not observe this pattern in our study, as germination seemed as plastic as other phenological traits.

Within each common garden, accessions from the Asian cluster flowered earlier than accessions from the other clusters, a phenological characteristic that might be an adaptation to Asia: (i) as a response to avoid biotic competition (Orsucci *et al*., 2020), thus compensating being in the colonization front and its consequent genetic load; or (ii) as a response to a rather unpredictable environment such as Asia (e.g., typhoon, monsoon), as observed in *Arabidopsis thaliana* (Simpson & Dean, 2002; Roux *et al*., 2006), that is a phylogenetically close species (Beilstein *et al*., 2006). It can also be a side consequence of a higher sensibility to photoperiod, as observed in tropical species such as rice (*Oryza sativa*) and maize (*Zea mays*) since the photoperiod is the main cue to the alternation of dry and humid seasons in these climates (Roux *et al*., 2006). According to this last hypothesis, early flowering reflects a different sensitivity to photoperiod, not an advantage of early flowering *per se*. In addition, several studies showed that, in several species such as *Arabidopsis thaliana* (Le Corre *et al*., 2002; Komeda, 2004) or *Triticum monococcum* (Yan *et al*., 2003), late flowering seems to be the ancestral character: if it is also the case for *C. bursa-pastoris*, it is probable that the “early flowering” phenotype recently evolved during *C. bursa-pastoris* range expansion in Asia.

Our study also showed that the European cluster flowered later than the other clusters, causing a longer time span for vegetative growth, and partly explaining the significantly larger rosette. In the Alpine accessions of *C. bursa-pastoris*, overwintering in the rosette state is frequently observed (Neuffer, 1990): a large rosette could be an adaptation towards harsh winter conditions in Europe, though accessions with large rosette were not specifically from the Alps. In our study, correlation between the rosette size and the flowering time was of *r* = 0.52 (see also Figure 2). It is not surprising to see an association between flowering trends and rosette characteristics, as it has been shown previously that QTLs associated with these two traits are closely linked in *C. bursa-pastoris* (Linde *et al*., 2001); likewise, in *Arabidopsis thaliana*, rosette leaf number has been shown to be sensitive to stimuli of flowering, such as shading (Cookson & Granier, 2006) or length of photoperiod (Lewandowska *et al*., 2017).

## Conclusion

Despite the large scale of the current study, involving three common gardens on as many continents and a large number of accessions representative of the natural range of the shepherd’s purse, we only detected phenotypic plasticity and adaptation signatures, but not of local adaptation. Does this indicate that local adaptation did not contribute to the rapid colonization of *C. bursa-pastoris?* It is probably too early to conclude, but the present study suggests that understanding local adaptation in *C. bursa-pastoris* and other ruderal species that went through a rapid range expansion would likely require a combination of more targeted reciprocal transfer experiments as well as more small scale experiments.

## Acknowledgements

We thank the Swedish Research Council and the Erik Philip Sörensen Foundation (to ML), Uppsala University and EMBO Short-Term Fellowship fund (to AC), and the H2020 European consortium B4Est (to MT) for funding. This research was also funded by the National Natural Science Foundation of China (grant No. 31870353, to HRH). AS was supported by an NSERC CGS-M and an Ontario Graduate Scholarship. We thank Julia Dankanich, Josefine Stångberg, Johanna Nyström, Sara Kurland and Uriel Gélin for field assistance in Uppsala; Jean-Tristan Brandenburg for R scripting support; Lin-Lin Wu, Zhi-Bing Xie, Qiu-Ling Guan, Gui-Yu Lin for field assistance in Guangzhou; Viktor Mollov and Niroshini Epitawalage for field assistance in Toronto; and finally, we also thanks Benoit Pujol, Benoit Facon, Denis Faure for a their comments on the first versions on the manuscript.

## Author Contribution

AC, SG, SW and MT planned and designed the common gardens. AC, SG, SW and ML directed the work. AC, AS, HH, DK, and KH performed experiments (installing the common garden and phenotyping every accessions). AC, MT, MO and PM analyzed datasets. AC and MT wrote the article, with thorough help from MO and PM. AC, MT, AS and HH contributed equally.

## Data availability

The data that support the findings of this study are openly available in [repository name e.g “figshare”] at http://doi.org/[doi], reference number [reference number].

---

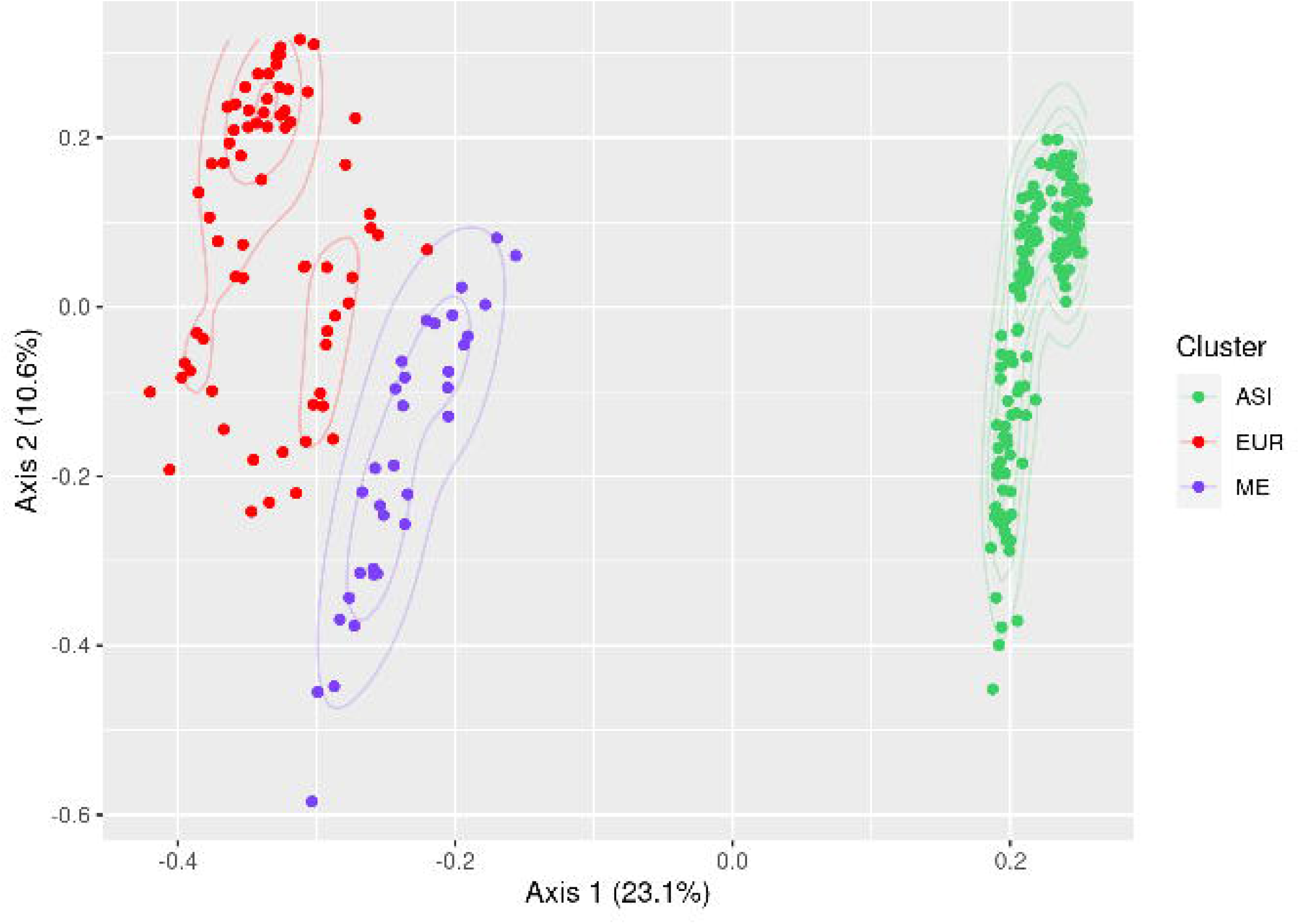

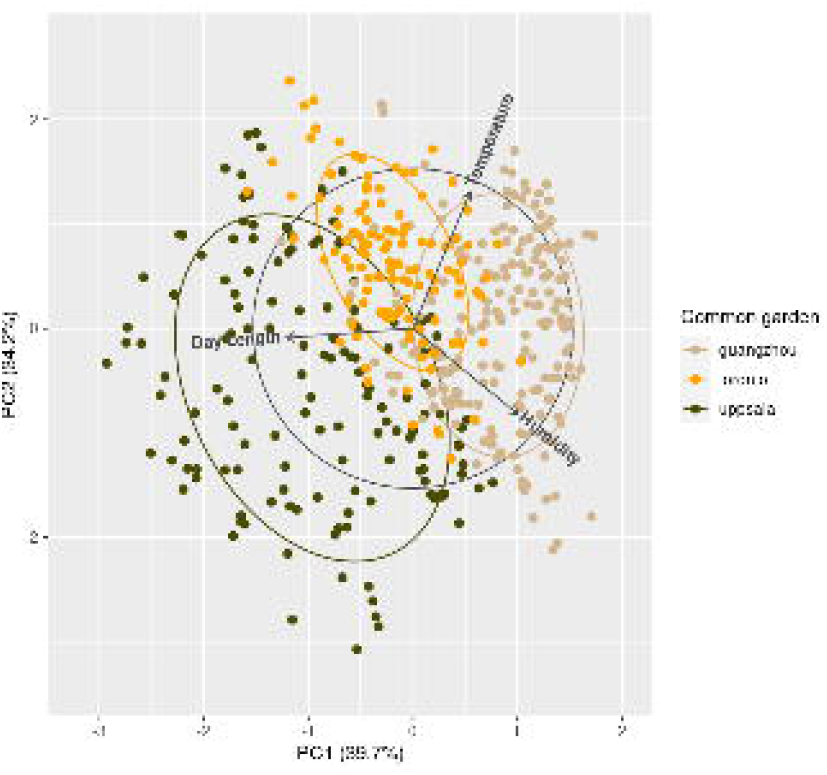

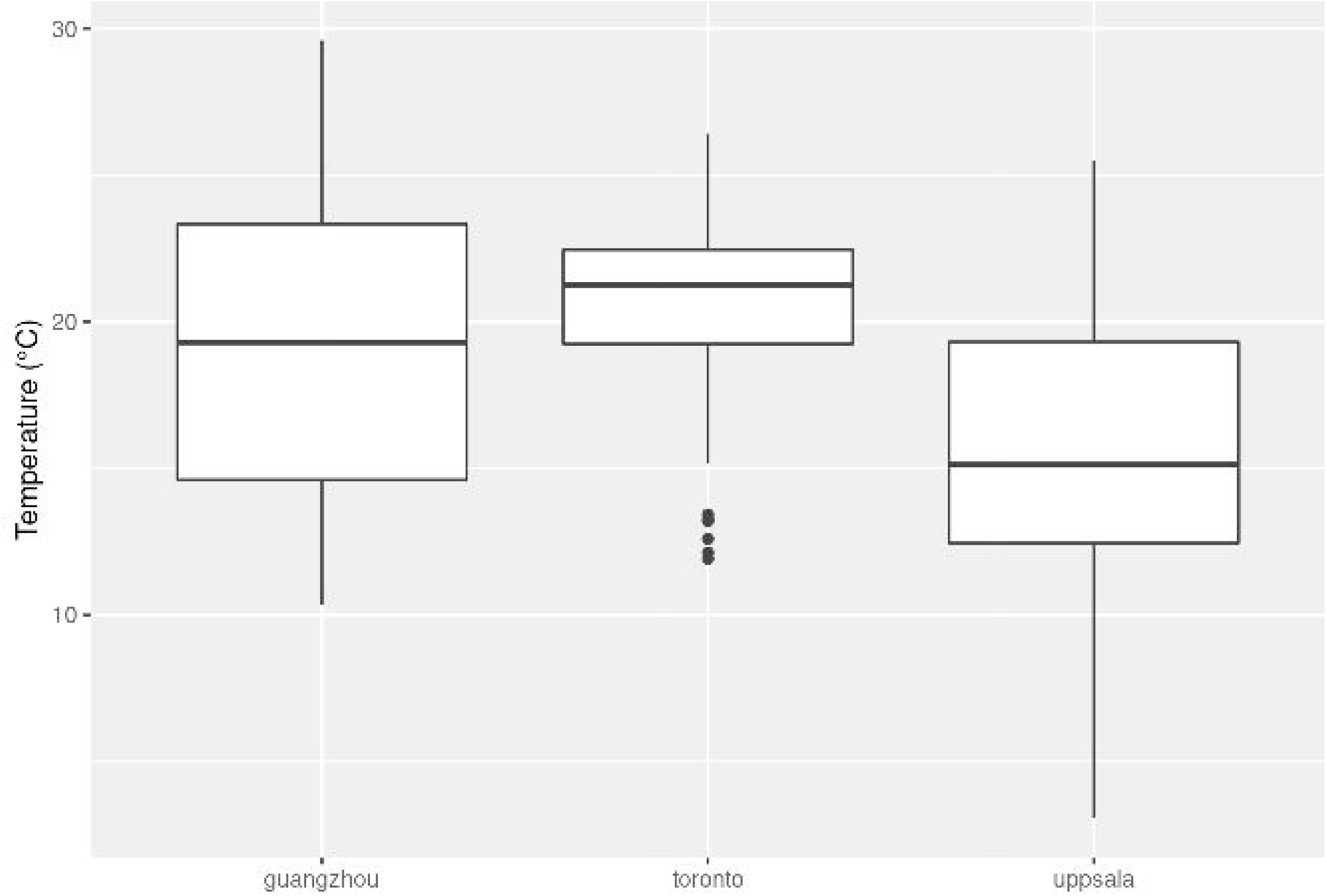

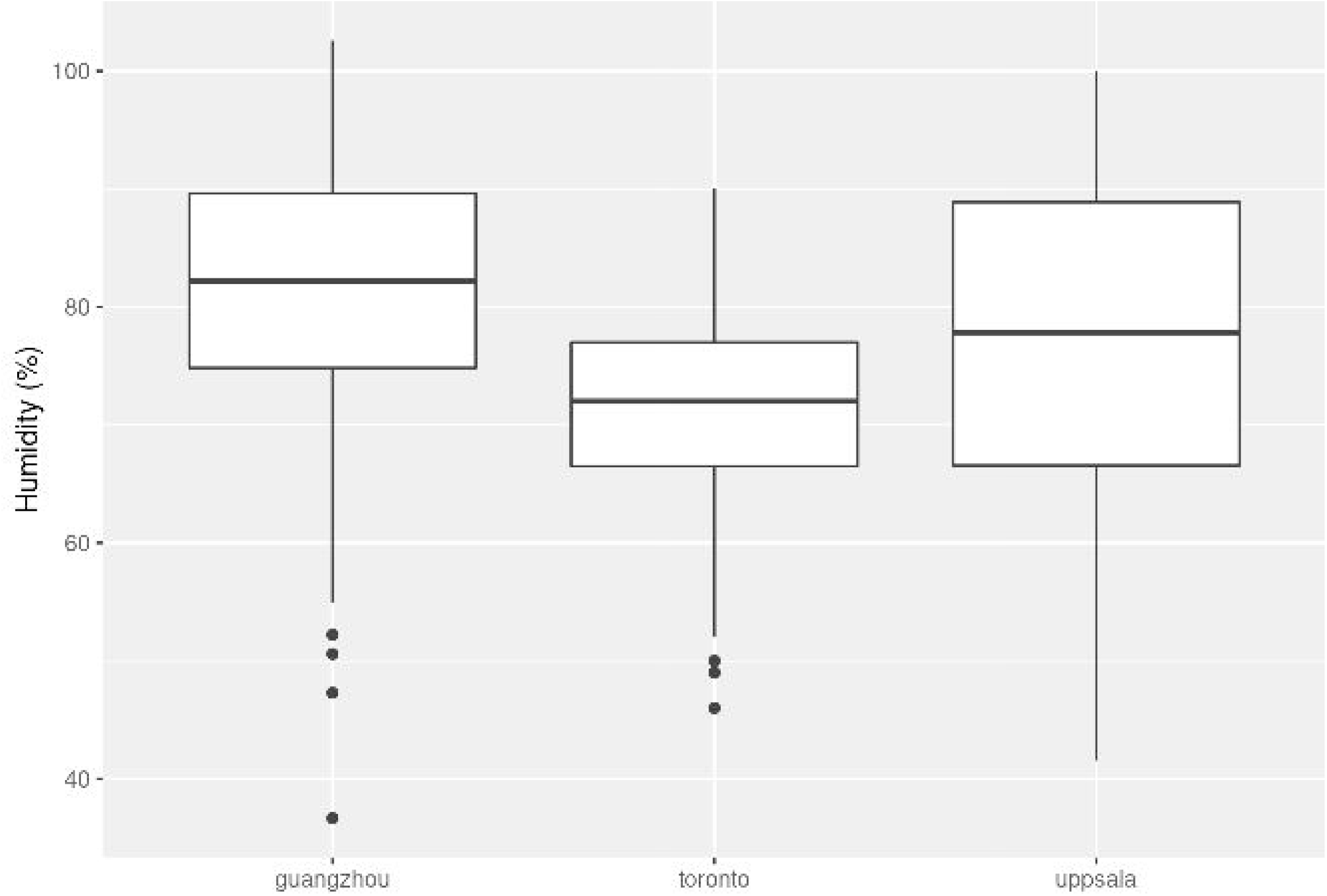

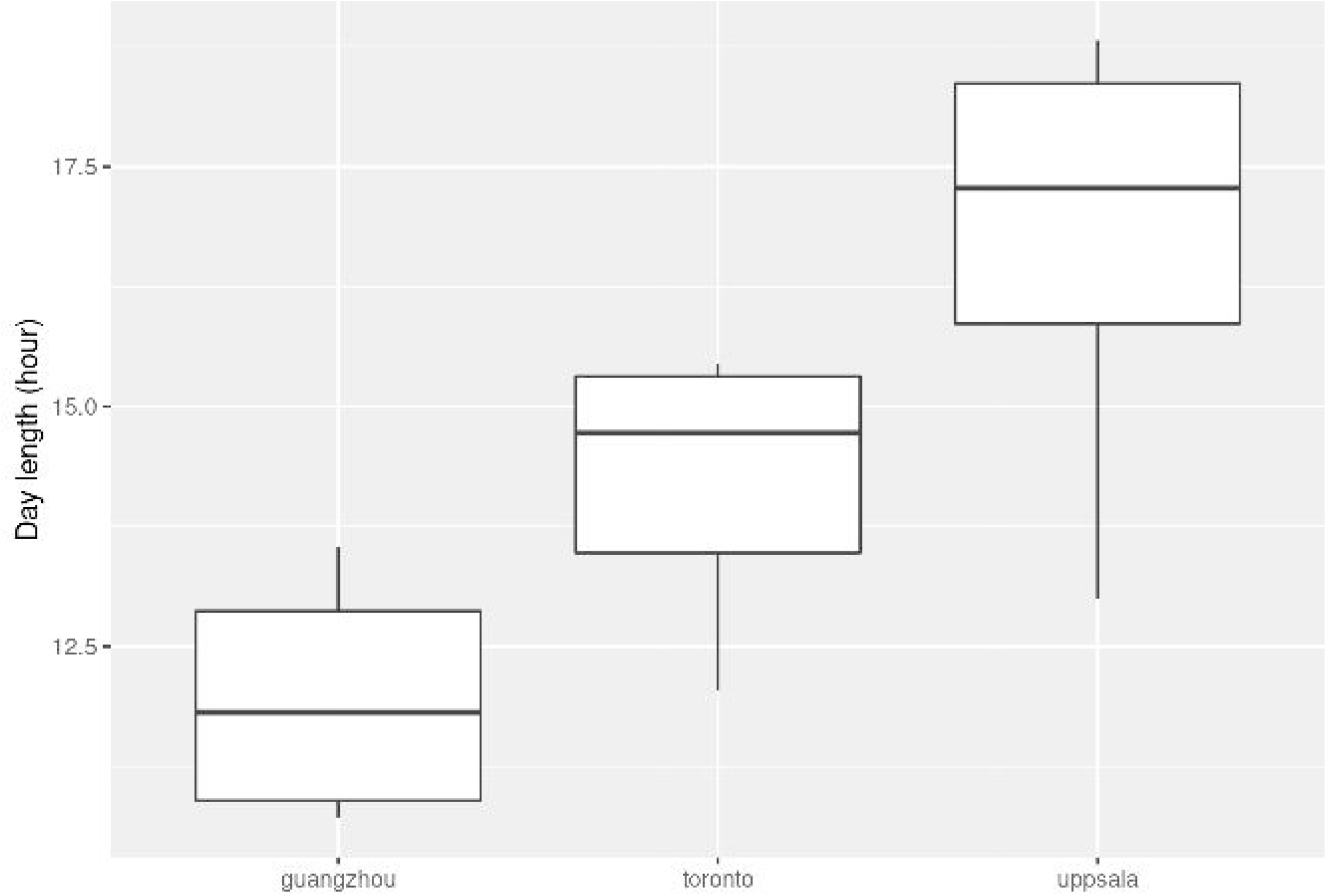

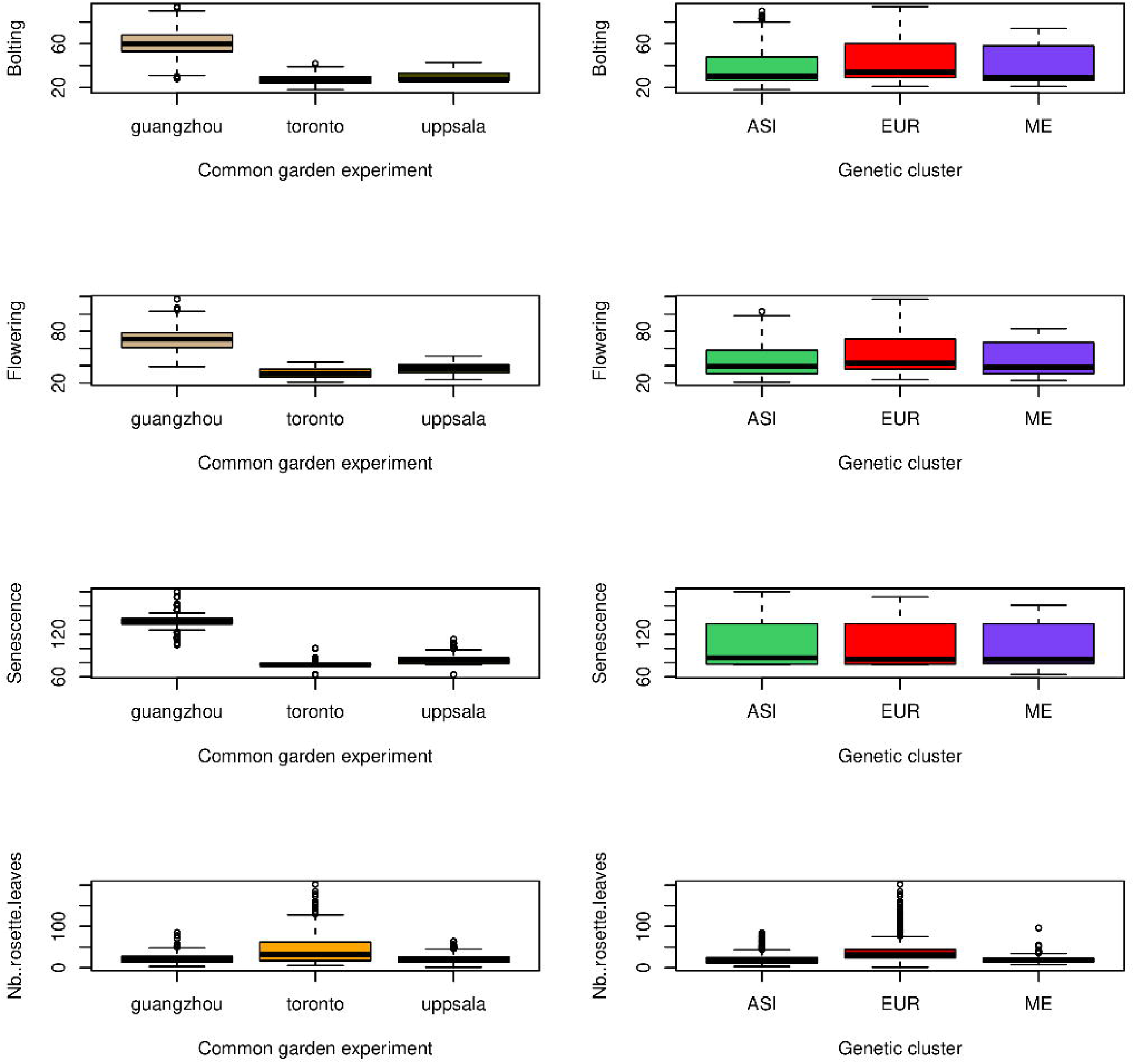

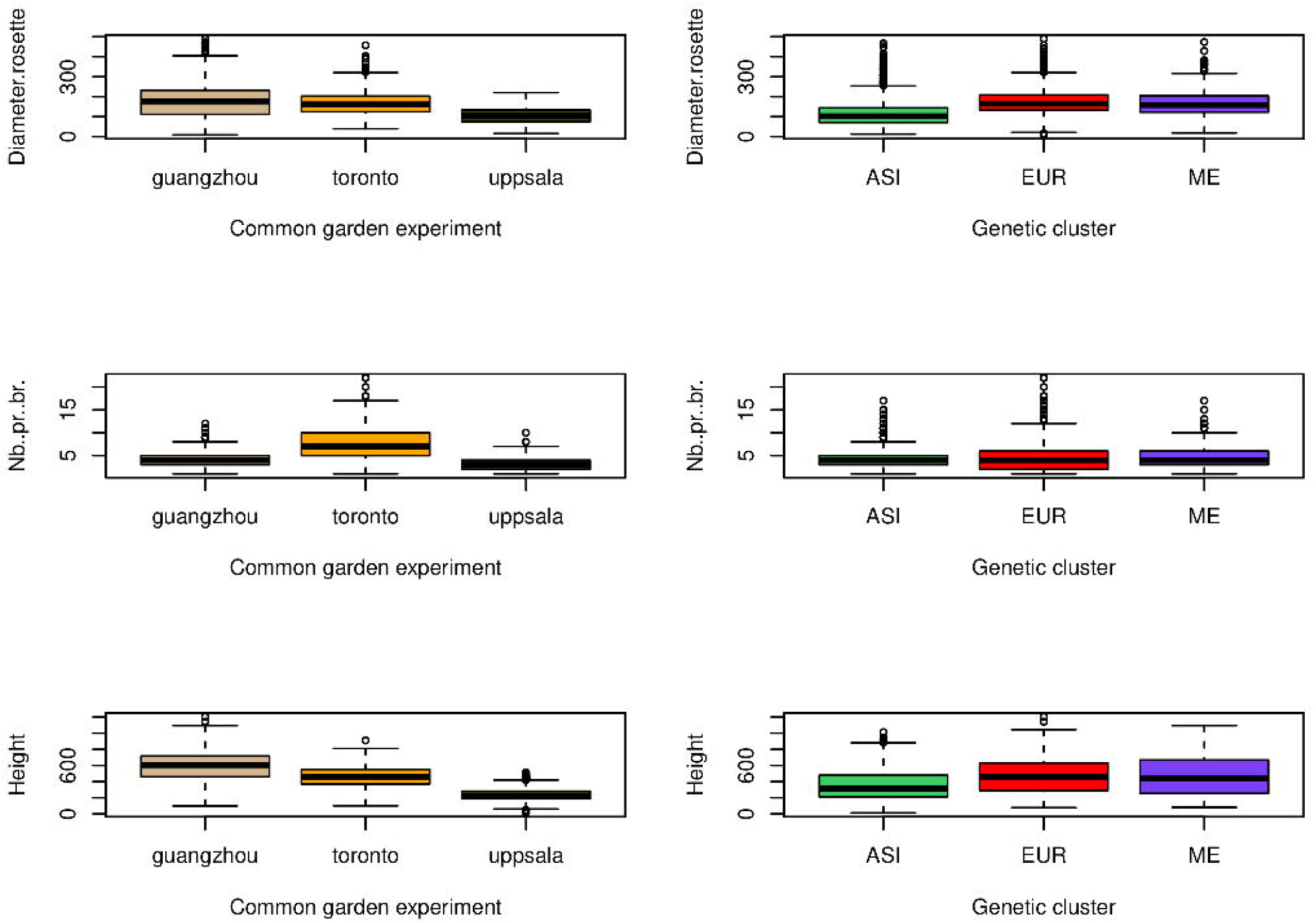

