## Supplementary file for "Phenotypic plasticity, population structure and adaptation in a young weed species with a worldwide distribution"

### Supplementary materials

**Preliminary analyses revealing the significance of common gardens and genetic clusters.** In order to study whether the interaction between common garden and genetic cluster was significant, we applied the following model to the global (and unbalanced) dataset:

Fitness*_ijkl_* ~ μ + cluster*_i_* + cg*_j_* + cluster*_i_* x cg*_j_* + accession*_k_* + block*_l_* + e*_ijkl_*	[S1],

where μ is the overall mean, ‘cluster’ and ‘cg’ are the fixed effects of respectively the genetic cluster and the common garden, ‘cluster x cg ’ is the interaction term, ‘block’ and ‘accession’ are uncorrelated random effects (‘accession’ being nested into ‘cluster’, and ‘block’ into ‘cg’), both normally distributed with its own mean and variance parameters, and ‘e’ is the residual. The distribution of the residual was fitted to a Normal distribution (function *lmer* of R package *lme4*; see Bates 2015).

For this preliminary analysis, we used the whole dataset, without balancing the block design: we wanted to assess whether the interaction was significant. Type II Wald chi-square test revealed that the interaction term was indeed significant for fitness (χ^2^ = 135.14, df = 4, *p* < 0.001, type II Wald chi-square test), justifying then the separation of the dataset (following the recommendation of Crawley, 2012). All values for other traits than fitness are summarized in Table 1.

### Tables

**Table S1.** List of the accessions used, along with the country where they were sampled and their genetic cluster (ASI, EUR, ME or none). The three next columns indicate whether the accessions died (0), survived but was removed from the dataset to balance the block design (X), or survived and were kept in the dataset (X*).

| accessionID | country | cluster | uppsala | guangzhou | toronto |
| --- | --- | --- | --- | --- | --- |
| 113 | China | ASI | X* | X* | 0 |
| 114 | China | ASI | X | X | X* |
| 115 | China | ASI | X* | X | X |
| 116 | China | ASI | X* | X | X* |
| 117 | China | ASI | X* | X | X |
| 118 | China | ASI | X* | X | X* |
| 119 | China | ASI | X* | X | X* |
| 120 | China | ASI | X* | 0 | X |
| 121 | China | ASI | X | 0 | X |
| 122 | China | ASI | X | X | 0 |
| 123 | China | ASI | X* | X | 0 |
| 125 | China | ASI | 0 | X | X |
| 126 | China | ASI | X* | X | X* |
| 127 | China | ASI | X* | X | 0 |
| 128 | China | ASI | X* | X* | 0 |
| 129 | China | ASI | X* | X | 0 |
| 130 | China | ASI | X* | 0 | 0 |
| 131 | China | ASI | X | X* | X* |
| 132 | China | ASI | X* | X | 0 |
| 133 | China | ASI | X* | 0 | 0 |
| 134 | China | ASI | X* | X | X* |
| 135 | China | ASI | 0 | X | 0 |
| 137 | China | ASI | X* | 0 | 0 |
| 139 | China | ASI | X* | X | 0 |
| 140 | China | ASI | X | X* | X* |
| 141 | China | ASI | X* | X | X* |
| 142 | China | ASI | X* | X* | 0 |
| 143 | China | ASI | X* | X* | 0 |
| 144 | China | ASI | X* | X* | 0 |
| 145 | China | ASI | X* | X | X* |
| 147 | China | ASI | X* | 0 | 0 |
| 148 | China | ASI | X | X | 0 |
| 149 | China | ASI | X | 0 | 0 |
| 150 | China | ASI | X* | X | X* |
| 151 | China | ASI | X* | X | X* |
| 152 | China | ASI | X | X | 0 |
| 153 | China | ASI | X* | 0 | 0 |
| 154 | China | ASI | 0 | 0 | X* |
| 155 | China | ASI | X* | X* | X* |
| 157 | China | ASI | X* | X* | 0 |
| 158 | China | ASI | X* | 0 | X* |
| 159 | China | ASI | X* | X | X* |
| 161 | China | ASI | X* | X | X* |
| 164 | China | ASI | X* | X* | 0 |
| 165 | China | ASI | X* | X | X |
| 166 | China | ASI | X* | X* | X* |
| 167 | China | ASI | X* | X | 0 |
| 169 | China | ASI | X* | X | 0 |
| 170 | China | ASI | X* | X | 0 |
| 171 | China | ASI | X | X | 0 |
| 175 | China | ASI | X | X | X* |
| 176 | China | ASI | X | X | 0 |
| 177 | China | ASI | X* | X* | 0 |
| 178 | China | ASI | X* | X | 0 |
| 179 | China | ASI | X* | 0 | 0 |
| 180 | China | ASI | X* | 0 | 0 |
| 181 | China | ASI | X* | X | 0 |
| 182 | China | ASI | X* | X | 0 |
| 183 | China | ASI | X | 0 | 0 |
| 185 | China | ASI | X | X | X* |
| 186 | China | ASI | X | X* | 0 |
| 187 | China | ASI | X | 0 | X |
| 188 | China | ASI | X | 0 | X* |
| 189 | China | ASI | X* | X* | 0 |
| 190 | China | ASI | X* | X* | 0 |
| 191 | China | ASI | X* | X* | 0 |
| 192 | China | ASI | X* | X* | 0 |
| 193 | China | ASI | X* | 0 | 0 |
| 195 | China | ASI | X* | X* | 0 |
| 196 | China | ASI | X* | X | 0 |
| 197 | China | ASI | X* | X* | 0 |
| 198 | China | ASI | X* | X | 0 |
| 199 | China | ASI | X* | 0 | X |
| 200 | China | ASI | X* | X | 0 |
| 201 | China | ASI | X* | X* | X* |
| 202 | China | ASI | X* | 0 | 0 |
| 203 | China | ASI | X* | X | X |
| 204 | China | ASI | X* | X | X* |
| 205 | China | ASI | X* | X | 0 |
| 206 | China | ASI | X* | X* | 0 |
| 207 | China | ASI | X* | X* | 0 |
| 208 | China | ASI | X* | X | 0 |
| 209 | China | ASI | X* | 0 | 0 |
| 210 | China | ASI | X* | 0 | 0 |
| 211 | China | ASI | X* | X | 0 |
| 212 | Taiwan | ASI | X* | 0 | X |
| 213 | China | ASI | X | X | X |
| 214 | China | ASI | X* | X* | 0 |
| 215 | China | ASI | X* | X | X* |
| 216 | China | ASI | X* | X | X* |
| 217 | China | ASI | X* | X | 0 |
| 218 | China | ASI | X* | X | X |
| 219 | China | ASI | X | X | X |
| 221 | China | ASI | X | 0 | X |
| 222 | China | ASI | 0 | 0 | X |
| 223 | China | ASI | X* | X* | X* |
| 224 | China | ASI | X | X | X* |
| 225 | China | ASI | X | 0 | 0 |
| 226 | China | ASI | X | X | X* |
| 228 | China | ASI | X* | X | X* |
| 231 | China | ASI | X* | 0 | X* |
| 232 | China | ASI | X* | X | X* |
| 233 | China | ASI | X* | X | X* |
| 234 | China | ASI | X* | X* | X* |
| 235 | China | ASI | X* | X | X* |
| 236 | China | ASI | X* | X | 0 |
| 237 | China | ASI | X | X | X |
| 238 | China | ASI | 0 | 0 | X* |
| 239 | China | ASI | X | X* | X |
| 240 | China | ASI | X | 0 | X |
| 242 | China | ASI | X* | X | X |
| 243 | China | ASI | X | 0 | X |
| 245 | China | ASI | X* | X | X* |
| 246 | China | ASI | X* | X* | X* |
| 247 | China | ASI | X* | X | 0 |
| 248 | China | ASI | X | X* | 0 |
| 249 | China | ASI | X* | X | 0 |
| 251 | China | ASI | X* | 0 | 0 |
| 252 | China | ASI | X* | X* | X* |
| 253 | China | ASI | X* | X | X* |
| 254 | China | ASI | X* | X | X* |
| 255 | China | ASI | X | X* | X* |
| 256 | China | ASI | X* | X* | X* |
| 257 | China | ASI | X* | X* | X* |
| 258 | China | ASI | X* | 0 | 0 |
| 259 | China | ASI | X | X* | X* |
| 260 | China | ASI | X* | X* | X* |
| 261 | China | ASI | X* | X* | X* |
| 262 | China | ASI | X* | 0 | X |
| 263 | China | ASI | X | X | X* |
| 264 | China | ASI | X* | X* | X |
| 265 | China | ASI | X* | X | 0 |
| 266 | China | ASI | X* | X | 0 |
| 267 | China | ASI | X | X* | 0 |
| 268 | China | ASI | X | X* | X* |
| 269 | China | ASI | X* | X* | X |
| 270 | China | ASI | X* | X* | X |
| 271 | China | ASI | X* | X | X |
| 272 | China | ASI | X* | X | X |
| 273 | China | ASI | X* | X* | X* |
| 1 | Italy | EUR | X* | X | X* |
| 10 | Spain | EUR | X | X | X* |
| 100 | Russia | EUR | X* | X | X* |
| 101 | Russia | EUR | X* | X | X* |
| 102 | Russia | EUR | X* | X* | X |
| 103 | Russia | EUR | X* | X | X* |
| 104 | Russia | EUR | X* | X | X* |
| 105 | Russia | EUR | X* | X* | X* |
| 106 | Russia | EUR | X* | X | X* |
| 11 | Spain | EUR | X | X* | X* |
| 12 | Spain | EUR | X* | X | X* |
| 13 | Spain | EUR | X* | X | X* |
| 14 | Spain | EUR | X* | X | X* |
| 15 | France | EUR | X* | 0 | 0 |
| 16 | France | EUR | X* | X | X |
| 17 | France | EUR | X* | X* | 0 |
| 18 | France | EUR | X* | X | 0 |
| 19 | France | EUR | X* | X | 0 |
| 2 | Italy | EUR | X* | X* | 0 |
| 20 | France | EUR | X* | X* | 0 |
| 21 | France | EUR | X* | X* | 0 |
| 22 | France | EUR | X* | X* | 0 |
| 25 | Russia | EUR | X | X | X* |
| 26 | Russia | EUR | X* | X | 0 |
| 27 | Russia | EUR | X* | X | 0 |
| 28 | Russia | EUR | X | X* | 0 |
| 29 | Russia | EUR | X* | X | 0 |
| 30 | Russia | EUR | X* | X | X* |
| 31 | Russia | EUR | X* | X | X* |
| 32 | Russia | EUR | X* | X* | X* |
| 33 | Russia | EUR | X | X | 0 |
| 34 | Russia | EUR | X* | X | X* |
| 35 | Czech | EUR | X* | X* | X* |
| 36 | France | EUR | X* | X* | X* |
| 37 | Greece | EUR | X | X | X* |
| 38 | Russia | EUR | X* | X | X* |
| 39 | Russia | EUR | X* | X | X* |
| 42 | Russia | EUR | X | X* | X |
| 43 | Russia | EUR | X* | X* | X* |
| 44 | Russia | EUR | X* | X* | X* |
| 45 | Russia | EUR | X* | X | X* |
| 46 | Russia | EUR | X* | X | X* |
| 47 | Russia | EUR | X | X | 0 |
| 54 | Russia | EUR | X | X | X* |
| 55 | Russia | EUR | X* | X* | X* |
| 56 | Russia | EUR | X* | X | X* |
| 57 | Russia | EUR | X* | X* | X* |
| 58 | Russia | EUR | X | X* | X* |
| 59 | Russia | EUR | X* | X | 0 |
| 60 | Russia | EUR | X* | X | 0 |
| 61 | Russia | EUR | X* | X* | X* |
| 62 | Russia | EUR | X* | X | X* |
| 63 | Russia | EUR | X* | X | X |
| 65 | Russia | EUR | X* | 0 | X* |
| 72 | Sweden | EUR | X | X | X* |
| 73 | Sweden | EUR | X | X* | X* |
| 74 | Sweden | EUR | X* | X | X* |
| 76 | United_Kingdom | EUR | X* | X | X* |
| 77 | United_Kingdom | EUR | X* | X | X* |
| 78 | United_Kingdom | EUR | X* | X | X* |
| 79 | France | EUR | X* | X | X* |
| 80 | France | EUR | X | X | X* |
| 87 | United_Kingdom | EUR | X* | X | X* |
| 88 | United_Kingdom | EUR | X | X | X* |
| 9 | Spain | EUR | X* | X | X* |
| 96 | United_States | EUR | X* | X | X* |
| 97 | Russia | EUR | X | X* | X* |
| 98 | Russia | EUR | X* | X* | X |
| 99 | Russia | EUR | X | X* | X* |
| 107 | United_States | ME | X* | X* | X* |
| 108 | United_States | ME | X* | X* | X* |
| 109 | United_States | ME | X* | X* | X* |
| 110 | United_States | ME | X* | X* | 0 |
| 111 | United_States | ME | X* | X* | 0 |
| 112 | United_States | ME | X* | X* | X* |
| 23 | Algeria | ME | X* | X* | X* |
| 24 | Algeria | ME | X* | X | X* |
| 3 | Italy | ME | X* | X* | X* |
| 4 | Italy | ME | X* | X* | 0 |
| 40 | Jordan | ME | X* | X* | X* |
| 41 | Jordan | ME | X* | X | X* |
| 49 | Israel | ME | X* | X* | 0 |
| 5 | Italy | ME | X* | X | X* |
| 50 | Israel | ME | X* | X | 0 |
| 51 | Israel | ME | X* | X* | 0 |
| 52 | Israel | ME | X* | X | 0 |
| 53 | Israel | ME | X* | X | 0 |
| 6 | Italy | ME | X | X* | 0 |
| 7 | Italy | ME | X | X | 0 |
| 8 | Italy | ME | X* | X | 0 |
| 81 | Syria | ME | X | X* | X* |
| 82 | Syria | ME | X* | X* | X* |
| 83 | Syria | ME | X* | X* | X* |
| 84 | Syria | ME | X* | X* | X* |
| 85 | Syria | ME | X* | X* | X* |
| 86 | Syria | ME | X* | X* | X* |
| 90 | Turkey | ME | X* | X | X* |
| 91 | Turkey | ME | X* | X | X* |
| 92 | Turkey | ME | X* | X* | X* |
| 93 | Turkey | ME | X* | X | X |
| 94 | Turkey | ME | X* | X* | X |
| 95 | United_States | ME | X* | X* | X |
| 124 | China |  | 0 | X | 0 |
| 136 | China |  | 0 | X | 0 |
| 138 | China |  | X | X | X |
| 146 | China |  | X | 0 | 0 |
| 160 | China |  | X | X | 0 |
| 172 | China |  | X | X | 0 |
| 173 | China |  | X | X | 0 |
| 184 | China |  | X | X | 0 |
| 184bis | China |  | 0 | X | 0 |
| 194 | China |  | X | X | 0 |
| 227 | China |  | 0 | X | X |
| 229 | China |  | X | X | 0 |
| 244 | China |  | X | 0 | X |
| 250 | China |  | X | X | X |
| 274 | China |  | X | 0 | 0 |
| 48 | Israel |  | 0 | X | X |
| 64 | Russia |  | X | X | X |
| 66 | Bosnia |  | X | X | X |
| 67 | Bosnia |  | X | X | 0 |
| 68 | Bosnia |  | X | X | 0 |
| 69 | Bosnia |  | X | X | 0 |
| 70 | Bosnia |  | X | X | X |
| 71 | Bosnia |  | X | X | 0 |
| 75 | Sweden |  | X | X | X |
| 89 | United_Kingdom |  | X | X | X |

**Table S2.** Average values of temperature, humidity and length of [photoperiod](https://www.sciencedirect.com/topics/earth-and-planetary-sciences/photoperiod) at Uppsala and for each month. The first and last month indicate the last date of the experiment.

| **Uppsala** | **2014** | | | | |
| --- | --- | --- | --- | --- | --- |
|  | **May (> 1st)** | **Jun.** | **Jul.** | **Aug.** | **Sep. (< 13th)** |
| **Temperature (°C)** | 10.7 | 13.4 | 19.5 | 17.7 | 14.2 |
| **Humidity (%)** | 72.0 | 73.9 | 73.9 | 82.2 | 86.3 |
| **Photoperiod (h)** | 16.9 | 18.6 | 18.1 | 15.8 | 13.7 |

**Table S3.** Average values of temperature, humidity and length of [photoperiod](https://www.sciencedirect.com/topics/earth-and-planetary-sciences/photoperiod) at Uppsala and for each month. The first and last month indicate the last date of the experiment.

| **Guangzhou** | **2014** | | **2015** | | | | |
| --- | --- | --- | --- | --- | --- | --- | --- |
|  | **Nov. (> 1st)** | **Dec.** | **Jan.** | **Feb.** | **Mar.** | **Apr.** | **May (< 12th)** |
| **Temperature (°C)** | 20.4 | 14.4 | 14.6 | 17.8 | 21.7 | 23.5 | 26.8 |
| **Humidity (%)** | 76.6 | 74.6 | 78.1 | 84.2 | 88.5 | 82.6 | 86.3 |
| **Photoperiod (h)** | 10.7 | 10.9 | 11.4 | 12.0 | 12.7 | 13.2 | 13.5 |

**Table S4.** Average values of temperature, humidity and length of photoperiod at Uppsala and for each month. The first and last month indicate the last date of the experiment.

| **Toronto** | **2014** | | | |
| --- | --- | --- | --- | --- |
|  | **Jun. (> 2nd)** | **Jul.** | **Aug.** | **Sep. (< 27th)** |
| **Temperature (°C)** | 21.3 | 21.5 | 21.7 | 18.3 |
| **Humidity (%)** | 72.4 | 69.1 | 69.7 | 74.1 |
| **Photoperiod (h)** | 15.4 | 15.1 | 14.1 | 12.7 |

**Table S5.** Predicted phenology timings (in days) at the common garden of Uppsala (U), Guangzhou (G) or Toronto (T), for each cluster (ASI, EUR and ME), using the GLMM model. Germination time at Toronto was not monitored.

| **Cluster** | **Germination** | | | **Bolting** | | | **Flowering** | | | **Senescence** | | |
| --- | --- | --- | --- | --- | --- | --- | --- | --- | --- | --- | --- | --- |
|  | **U** | **G** | **T** | **U** | **G** | **T** | **U** | **G** | **T** | **U** | **G** | **T** |
| ASI | 4.0 | 2.1 | - | 27.6 | 56.9 | 27.3 | 34.3 | 66.9 | 31.0 | 85.0 | 138.3 | 77.6 |
| EUR | 2.4 | 0.6 | - | 30.4 | 65.3 | 29.8 | 39.5 | 76.8 | 34.7 | 84.1 | 139.4 | 79.0 |
| ME | 1.6 | 0.2 | - | 27.6 | 60.3 | 24.6 | 34.3 | 70.2 | 27.7 | 82.7 | 136.5 | 77.5 |

**Table S6.** Tests of the parameters of Finlay Wilkinson Regression (the FWR model) for the phenological traits: bolting time (BT), flowering time (FT), senescence time (ST), and flowering time span (SP). Each cell is structured as x/y, x being the significance of the pairwise mean difference (Welch’s t-test), and y the significance of the pairwise variance difference (Fisher’s F-test). Significance levels are: *p**** < 0.001; *p** <* 0.01; *p** < 0.05; *p^n.s.^* > 0.05.

| **Hypothesis** | **Intercept** | | | | | **Slope** | | | | |
| --- | --- | --- | --- | --- | --- | --- | --- | --- | --- | --- |
|  | **Fitness** | **BT** | **FT** | **ST** | **SP** | **Fitness** | **BT** | **FT** | **ST** | **SP** |
| ASI ≠ EUR | n.s. / n.s. | n.s. / n.s. | n.s. / n.s. | n.s. / n.s. | n.s. / n.s. | n.s. / n.s. | * / n.s. | n.s. / n.s. | n.s. / n.s. | n.s. / n.s. |
| ASI ≠ ME | n.s. / n.s. | *** / *** | n.s. / *** | n.s. / n.s. | n.s. / *** | ** / n.s. | n.s. / *** | n.s. / *** | n.s. / n.s. | n.s. / *** |
| EUR ≠ ME | n.s. / n.s. | * / *** | n.s. / *** | n.s. / n.s. | n.s. / *** | ** / n.s. | n.s. / *** | n.s. / *** | n.s. / n.s. | n.s. / *** |

**Table S7**. Variance ratio of fitness, for different genetic clusters (ASI, EUR and ME), using the whole dataset: statistics, p-values and degree of freedom (Df) of the Fisher’s F-test on the unbalanced dataset. Significance levels are : *p**** < 0.001; *p** <* 0.01; *p** < 0.05; *p^n.s.^* > 0.05.

| **Hypothesis** | **Estimate** | **Df (numerator/denominator)** |
| --- | --- | --- |
| ASI/EUR = 1 | 0.80*** | 1422/905 |
| ME/ASI = 1 | 2.23*** | 461/1422 |
| ME/EUR = 1 | 1.79*** | 461/905 |

### Figures

**Figure S1.** Multi-Dimensional Scaling of the GBS dataset (first two components, respectively explaining 23.1% and 10.6% of the genetic variance), clustering the accessions into three genetic clusters: ASI (in green), EUR (in red), and ME (in blue).

**Figure S2.** Principal component analysis (PCA) of the environmental data: day length, temperature and humidity. Dots correspond to daily records: 139 records for Uppsala (brown), 193 for Guangzhou (beige) and 118 for Toronto (orange).

**Figure S3.** Box plot of the temperature (in degree Celsius) according to different common garden.

**Figure S4.** Box plot of the humidity (in percentage) according to different common garden.

**Figure S5.** Box plot of the day length (in hour) according to different common garden.

**Figure S6.** Boxplots of phenotypic and phenological traits using the whole dataset: Bolting time, Flowering time, Senescence time, Number of rosette leaves, Diameter of rosette, Number of primary branches, and height of the inflorescence. Left column is according to the common garden (Uppsala in brown, Toronto in orange, and Guangzhou in beige), and right columns is according to the genetic cluster (ASI in green, EUR in red, and ME in blue).
